# Sequence-specific and programmable catalytic degradation of extracellular microRNAs by membrane-anchored DNA nanomachines

**DOI:** 10.64898/2026.09.30.755580

**Authors:** Ziyan Zhou, Linghao Zhang, Xiaowei Wang, Chunyi Zhang, Qiming Yan, Shihui Wang, Huiyu Liu, Xin Su

## Abstract

Extracellular microRNAs (miRNAs) contribute to metastasis through communication between tumour cells and their microenvironment, yet their sustained, sequence-selective depletion remains challenging. Here we report Programmable RNA-Intercepting Molecular Eraser (PRIME), a membrane-anchored DNA nanomachine that organizes miRNA recognition and catalytic degradation at the cell surface. A double-tetrahedral DNA scaffold positions a programmable recognition hairpin beside RNase H. Target binding generates an RNA/DNA hybrid for RNase H-mediated cleavage, and product release resets the hairpin for subsequent degradation cycles. Membrane-anchored PRIME depleted matched extracellular miRNAs by 70.6– 89.4% across lung, breast and cervical cancer cell models. Multi-target PRIME (M-PRIME) inhibited migration by 71.3–96.0% in Transwell assays across these models. To evaluate its therapeutic potential, we established an orthotopic lung-tumour model and an early-intervention breast-cancer lung-colonization model in mice. Aptamer-directed M-PRIME combined with gemcitabine reduced pulmonary tumour burden by 74.5% in established lung tumours and 93.2% in the lung-colonization model. These effects were consistent with PRIME-mediated extracellular miRNA degradation, with circulating target miRNA levels reduced by 77.5–98.7% relative to gemcitabine alone. Together, these findings establish a programmable material strategy for sustained extracellular RNA depletion and support its potential to enhance chemotherapy in preclinical tumour models.

## Introduction

Metastasis is not solely a cell-autonomous consequence of tumour evolution; it is a multicompartment process coordinated by the exchange of signals among tumour cells, stromal and immune populations, vascular endothelium and distant-organ niches^1,2,3^. Extracellular microRNAs (miRNAs) offer sequence-defined targets for intervention^4^ because they can carry regulatory information between cells and alter gene expression in recipient cells^5,6^. Carried by extracellular vesicles^7^, lipoproteins^8^ or RNA-binding-protein complexes^9^, miRNAs can persist outside cells and remodel gene networks controlling vascular permeability, cell motility, survival, metabolic adaptation and pre-metastatic niche formation^10,11^. Tumour-derived miR-105^12^, miR-122^13^ and miR-181c^14^, for example, have been associated with endothelial-barrier disruption, metabolic reprogramming and organ-selective metastatic colonization. Extracellular miRNAs can thus contribute to the effects of tumours on neighbouring and distant tissues, in addition to serving as circulating biomarkers. This extracellular mode of action also creates a distinct therapeutic challenge: pathogenic miRNAs are continuously secreted, dispersed at low local concentrations and redistributed among heterogeneous recipient cells. Effective intervention therefore focuses on eliminating defined miRNA inputs at the sites of signal release and exchange, before their uptake by recipient cells and the onset of downstream responses.

Most miRNA-directed technologies were developed to alter intracellular miRNA activity. Antagomirs, chemically modified antisense oligonucleotides^15^, miRNA mimics, sponges^16^ and RNA-editing or RNA-cleavage platforms^17^ can inhibit, replace or destroy miRNAs after entering target cells^18,19,20^. Their efficacy, however, remains coupled to tissue distribution, cellular uptake, endosomal escape, intracellular stability and the attainment of sufficient local stoichiometry. More fundamentally, intracellular intervention does not directly remove the extracellular signals that connect different cellular compartments: even efficient inhibition in a subset of tumour cells leaves secreted miRNAs available to act on stromal, endothelial, immune or distant recipient cells. Strategies that inhibit extracellular-vesicle release or uptake, capture vesicles or scavenge total extracellular RNAs shift the intervention outside the cell, but can perturb broader transport pathways or lack sequence specificity; capture-based approaches are also limited by finite binding capacity^21^. These limitations motivate approaches that selectively and sustainably remove extracellular miRNAs. Intercepting metastatic communication calls for a material that remains at the extracellular cell boundary, recognizes selected RNA sequences and catalytically consumes accessible target molecules. The capacity to accommodate multiple miRNA inputs is also central to this design, as metastatic signalling involves regulatory networks regulated by multiple extracellular species.

DNA nanotechnology provides a route to such an interface because molecular recognition, reaction geometry and biological localization can be encoded within one addressable material^22,23,24^. Base-pairing programmability enables recognition domains, catalytic components and targeting ligands to be positioned with nanoscale control^25,26,27^, while DNA tetrahedra^28^, origami frameworks^23,24,25^ and dynamic nanomachines^29,30^ provide defined architectures for presenting ligands, organizing enzymes, sensing nucleic acids and engaging cellular membranes^28^. Enzyme-coupled nucleic-acid reactions^31^ combine sequence-specific recognition with enzymatic cleavage, enabling the selective degradation of nucleic-acid targets^32,33,34^. We therefore hypothesized that a DNA-based nanomaterial could position RNase H close to a miRNA-recognition unit and retain both at the cell surface to support repeated extracellular miRNA degradation.

Here we developed Programmable RNA-Intercepting Molecular Eraser (PRIME), a membrane-anchored DNA nanomachine for selective, sustained degradation of extracellular miRNAs implicated in metastatic communication (Fig. 1). PRIME integrates an exchangeable miRNA-recognition hairpin and DNA-tethered RNase H on a double-tetrahedral scaffold, with cholesterol or tumour-cell-binding aptamers supporting membrane anchoring. Target binding forms an RNA/DNA hybrid for RNase H-mediated cleavage, and product release resets the hairpin for subsequent degradation cycles. PRIME showed sequence-specific responses to repeated miRNA inputs in solution and on living-cell membranes. To guide target selection, we integrated miRNA abundance in TCGA tumour profiles, evidence of extracellular occurrence in GEO datasets and literature-supported associations with migration or metastasis. Membrane-anchored PRIME depleted matched extracellular miRNAs by 70.6–89.4% across lung, breast and cervical cancer cell models. Combining three target-specific constructs yielded multi-target PRIME (M-PRIME), which inhibited migration by 71.3–96.0% in Transwell assays across these models. For *in vivo* evaluation, we established an orthotopic H1299-Luc lung-tumour model and an early-intervention MDA-MB-231-Luc breast-cancer lung-colonization model. Aptamer-directed M-PRIME combined with gemcitabine reduced pulmonary tumour burden by 74.5% in established lung tumours and 93.2% in the lung-colonization model. Circulating target miRNAs declined by 77.5–98.7% relative to gemcitabine alone, supporting a contribution of PRIME-mediated extracellular miRNA degradation to these therapeutic effects. These results define a cell-surface catalytic approach to controlling extracellular RNA signals and provide a basis for combining targeted miRNA depletion with chemotherapy.

**Figure 1.**
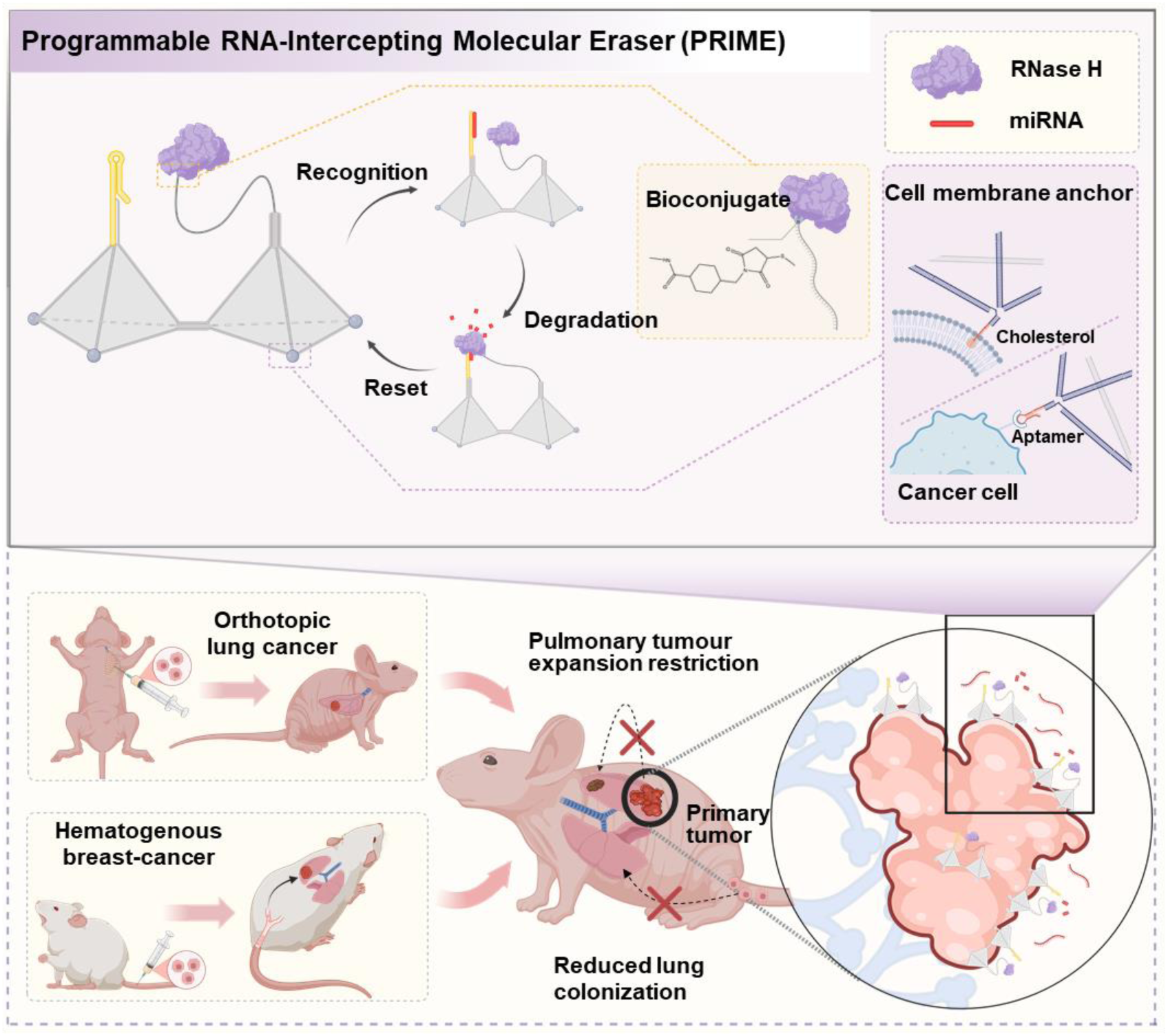
PRIME comprises a double-tetrahedral DNA scaffold that co-localizes a hairpin miRNA-recognition unit with an RNase H–DNA catalytic unit. Cholesterol modification anchors PRIME to the plasma membrane in cultured-cell experiments, whereas a tumour-cell-binding aptamer supports targeted membrane association *in vivo*. A cognate extracellular miRNA opens the hairpin and forms a local RNA/DNA hybrid, within which RNase H selectively cleaves the RNA strand. Dissociation of the cleavage products restores the recognition unit for subsequent degradation cycles. Exchanging the recognition hairpin changes the target of an individual PRIME construct; mixing multiple target-specific constructs yields M-PRIME which is evaluated for the enhancement of gemcitabine response in established orthotopic lung tumours and during early breast-cancer lung colonization.

## Results

### Construction and structural characterization of PRIME

PRIME was engineered as a modular DNA–enzyme nanostructure that spatially organizes miRNA recognition and catalytic degradation on adjacent tetrahedral units. The recognition tetrahedron displayed a hairpin miRNA-binding unit, whereas the catalytic tetrahedron carried a docking strand for sequence-specific hybridization of the covalent RNase H–DNA conjugate (Fig. 2a and Supplementary Fig. 1). The two functional units were assembled through programmed hybridization to form the double-tetrahedral PRIME scaffold. To prepare the catalytic unit, RNase H was functionalized with Sulfo-SMCC and covalently coupled to thiolated SH-DNA, allowing the resulting RNase H–DNA conjugate to be incorporated into the scaffold through sequence-specific hybridization. Native PAGE resolved the progressive assembly of the individual tetrahedra, functionalized subunits and complete PRIME structure (Supplementary Fig. 2). MALDI–TOF mass spectrometry detected component peaks for RNase H and SH-DNA at m/z 18,820.5 and 19,087.7, respectively, together with a higher-mass conjugate-associated species at m/z 38,684.4 (Fig. 2b and Supplementary Fig. 3), supporting formation of the RNase H–DNA conjugate. AFM imaging revealed discrete nanoscale particles and an increase in apparent particle dimensions following enzyme loading (Fig. 2c). Together, these measurements support formation of the RNase H–DNA conjugate and assembly of the PRIME nanostructure.

**Figure 2.**
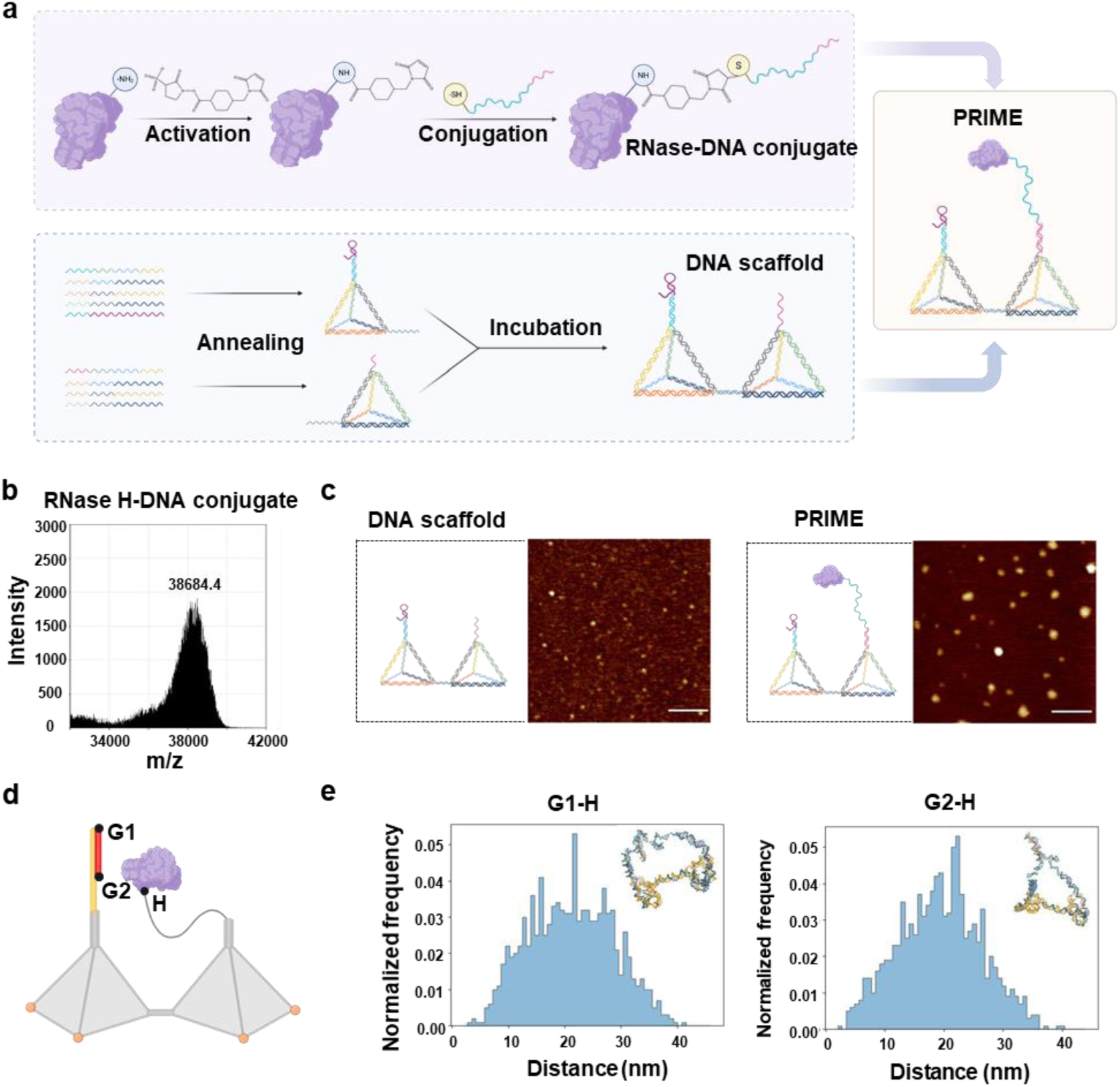
Construction and structural characterization of PRIME. a, Synthetic workflow for PRIME, including Sulfo-SMCC activation of RNase H, conjugation to thiol-modified SH-DNA, annealing of the two functional DNA tetrahedral units and hybridization-mediated loading of the RNase H-DNA conjugate. b, MALDI-TOF mass spectrum of the RNase H–DNA conjugate. c, Representative AFM images of the DNA scaffold of PRIME and enzyme-loaded PRIME. Scale bars, 200 nm. d, Reference positions used for distance measurements in the PRIME scaffold model. e, Simulated G1-H and G2-H distance distributions describing geometric sampling between the recognition region and the nucleotide position representing the RNase H-loading site.

Because efficient RNA cleavage requires physical access of RNase H to the captured substrate, we next examined whether the double-tetrahedral geometry supports productive enzyme–RNA encounters. Coarse-grained oxDNA modelling showed broad distributions of the distances between the substrate-recognition region (G1 and G2) and the RNase H-loading site (H), consistent with flexibility that could permit enzyme– substrate encounters (Fig. 2e).^35,36^ Complementary distance and root-mean-square fluctuation analyses indicated that the double-tetrahedral core preserves the spatial organization of the recognition and catalytic units while permitting local motion around the substrate-binding and enzyme-loading regions (Supplementary Figs. 4 and 5).^37^

### PRIME selectively degrades target miRNAs *in vitro*

To determine whether PRIME selectively recognizes and degrades target miRNAs, the reaction was monitored using fluorophore–quencher-labelled constructs. Each recognition unit was designed as a target-complementary stem-loop containing an exposed toehold, enabling cognate miRNA binding to initiate strand displacement while maintaining a closed state in the absence of target. In the closed hairpin state, Cy5 fluorescence was suppressed by BHQ3. Binding of a cognate miRNA opened the hairpin via toehold strand displacement^38^, restored fluorescence and generated a local RNA/DNA hybrid that acted as the substrate for RNase H (Fig. 3a). RNase H subsequently cleaved the RNA strand, allowing the degradation products to dissociate and the recognition unit to reset for subsequent cycles. After miR-21 addition, complete PRIME produced a transient fluorescence pulse, whereas RNase H-deficient PRIME showed sustained fluorescence and the no-target control remained near baseline (Fig. 3b). Native PAGE further confirmed that RNase H only cleaved RNA/DNA hybrids but not single-stranded RNA, single-stranded DNA or double-stranded DNA (Supplementary Fig. 6). This substrate requirement couples RNase H-mediated cleavage to formation of the RNA/DNA hybrid by the recognition unit.

**Figure 3.**
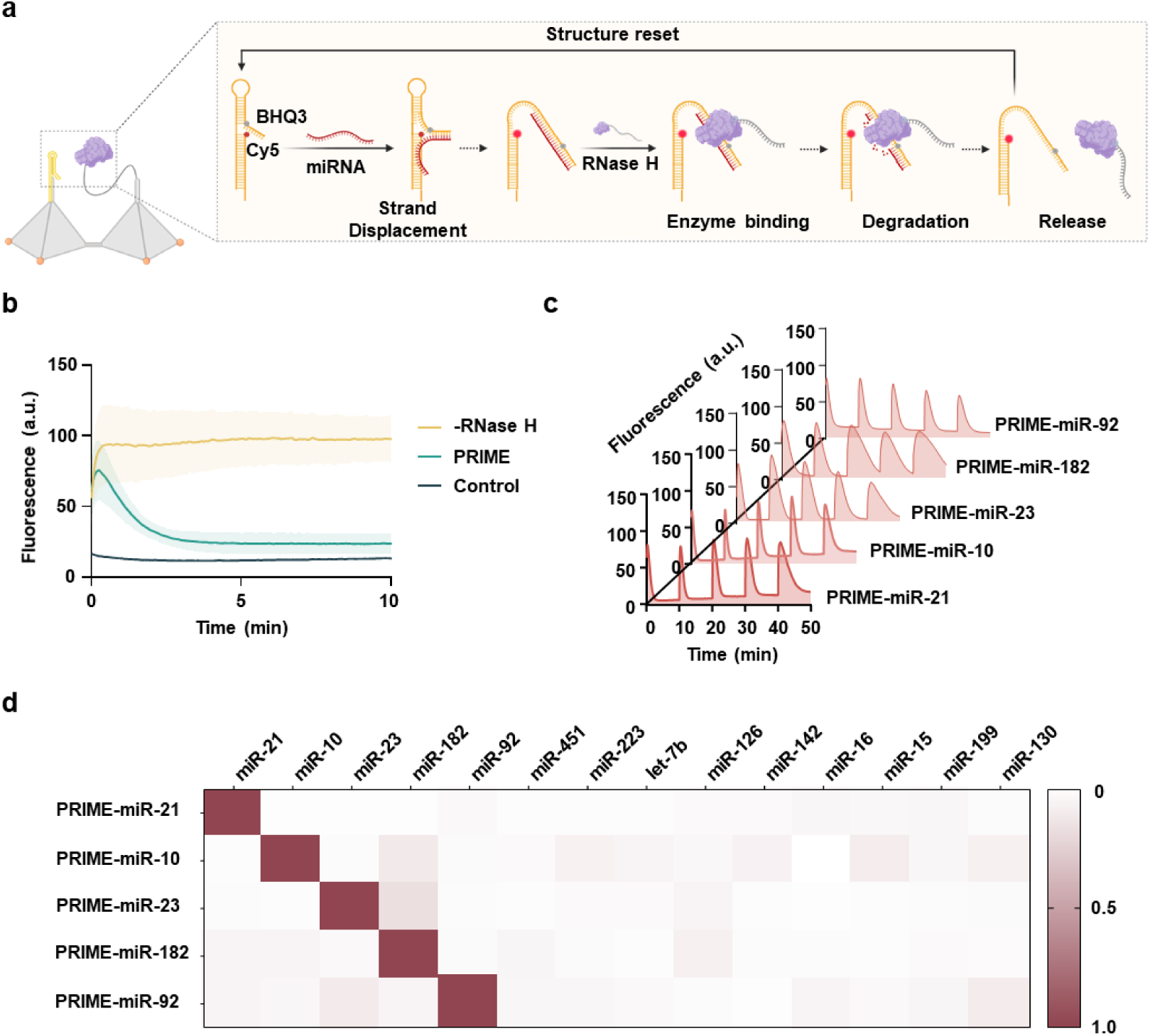
PRIME shows sustained and sequence-specific miRNA degradation *in vitro*. a, Reaction of fluorophore–quencher-labelled PRIME. A cognate miRNA opens the Cy5–BHQ3-labelled hairpin and forms an RNA/DNA hybrid. RNase H selectively cleaves the RNA strand, releasing the cleavage products and resetting the recognition unit for subsequent target binding. b, Real-time fluorescence profiles of complete PRIME, RNase H-deficient PRIME and the no-target control. Lines show the mean of three independent experiments, and shaded areas indicate the range. c, Repeated-input fluorescence-response assay for five target-specific PRIME constructs. Sequential additions of the corresponding cognate miRNA generate successive fluorescence pulses. Curves show the mean fluorescence response from three independent experiments. Data are mean ± s.d. from three independent experiments. d, Specificity matrix of fluorescence-trace AUC values for the five PRIME variants, normalized to the cognate-target response for each variant. PRIME and miRNA were used at 100 nM, and all reactions were performed at 37 °C.

To assess whether PRIME could sustain degradation, PRIMEs targeting miR-21, miR-10, miR-23, miR-182 and miR-92 were subjected to successive additions of their respective cognate miRNAs. Each cycle of miRNA input generated a fluorescence pulse (Fig. 3c). The response matrix showed strong signals for cognate PRIME–miRNA pairs and minimal responses to non-cognate miRNAs within the tested panel (Fig. 3d). These results support sustained and sequence-selective miRNA degradation of PRIME.

### Membrane-anchored PRIME depletes extracellular miRNAs on living cells

To determine whether PRIME could function as an extracellular catalytic interface on living cells, cholesterol-modified PRIME was anchored to the plasma membrane through hydrophobic insertion. Membrane loading was first quantified independently of catalytic activity using an unquenched Cy5-labelled construct. Cell-associated fluorescence increased with PRIME concentration and approached saturation at 300 nM, which was used for subsequent fluorescence imaging and flow-cytometry experiments (Supplementary Fig. 7). Native PAGE showed the structural stability of the PRIME nucleic-acid scaffold for 48 h in medium containing 10% fetal bovine serum (Supplementary Fig. 8). Fluorescence imaging and flow cytometry further showed persistent cell-associated signals over the same period (Supplementary Fig. 9).

Having established membrane localization, we next examined whether surface-anchored PRIME preserved the repeated target-recognition and degradation behaviour observed *in vitro*. Total internal reflection fluorescence (TIRF) microscopy was used to selectively monitor fluorophore–quencher-labelled PRIME within the membrane-proximal region. Sequential additions of miR-21 produced repeated increases in fluorescence followed by signal decay in the same cells, consistent with target-induced hairpin opening, formation of an RNA/DNA hybrid, RNase H-mediated RNA cleavage and restoration of the recognition unit (Fig. 4a and 4b). RNase H-deficient PRIME displayed target-dependent fluorescence activation but no signal decay, whereas cells not exposed to miR-21 remained near baseline (Supplementary Fig. 10). Flow cytometry reproduced these response patterns at the population level (Fig. 4c). These repeated responses indicate that membrane anchoring preserves the target-induced reaction cycles observed in solution.

**Figure 4.**
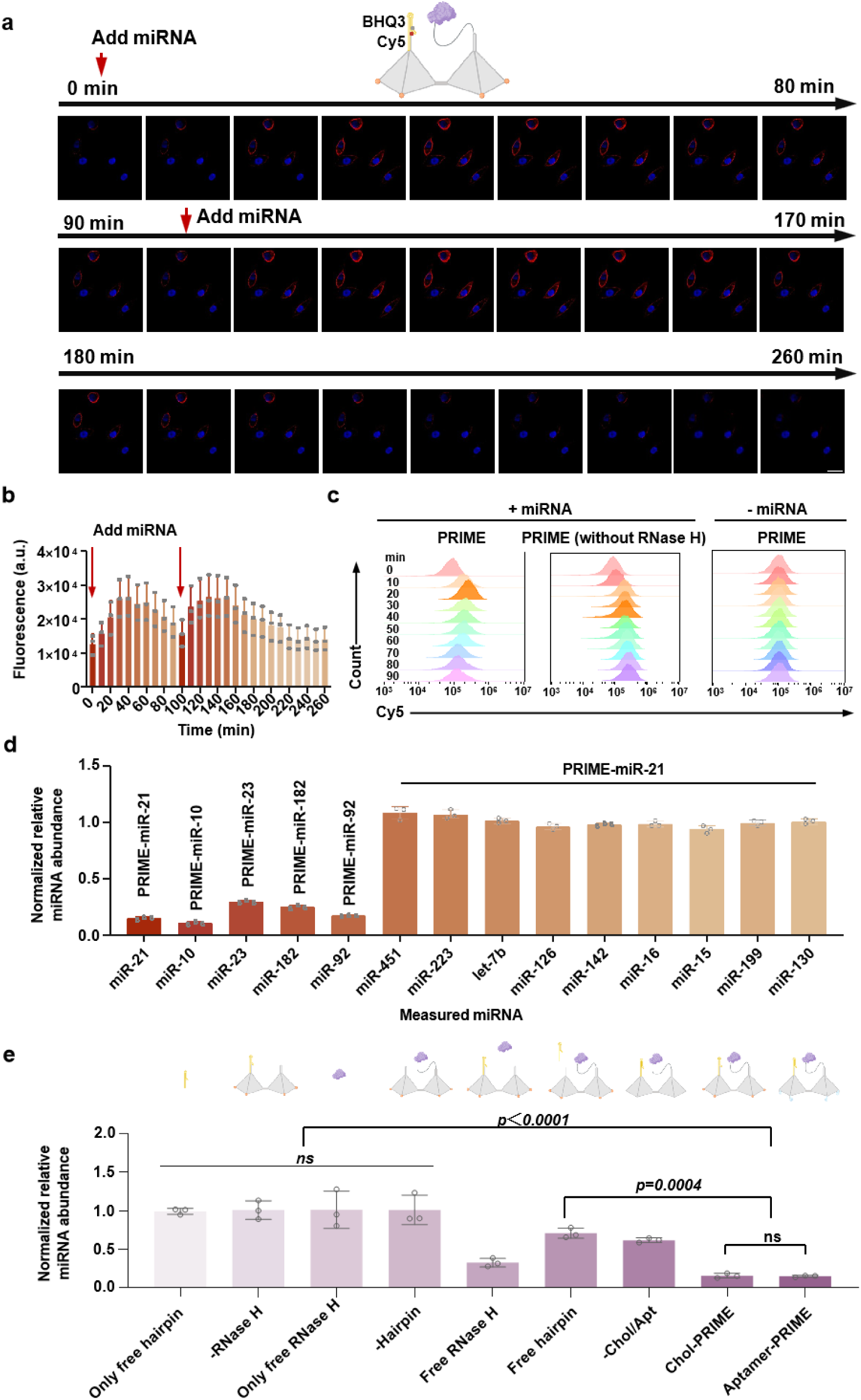
Membrane-anchored PRIME responds to repeated miRNA inputs and reduces extracellular miRNA abundance in cell cultures. a, Representative fluorescence images of A549 cells bearing fluorophore–quencher-labelled, cholesterol-anchored PRIME during sequential extracellular miR-21 exposure. Scale bar, 10 μm. b, Membrane-proximal fluorescence intensities in panel a, showing repeated target-induced activation of PRIME. Data are mean ± s.d. from three independent experiments. c, Flow-cytometry analysis of cell-associated reporter fluorescence in cells bearing complete PRIME, RNase H-deficient PRIME or complete PRIME without added miRNA. A total of 10,000 events were collected per sample. d, RT-qPCR quantification of extracellular miRNA abundance after 48 h treatment with matched or non-matched PRIME variants in A549 (miR-21, miR-10, miR-23, miR-451, miR-223, let-7b, miR-126, miR-142, miR-16, miR-15, miR-199 and miR-130), MDA-MB-231 (miR-182) and HeLa (miR-92) cells. e, RT–qPCR analysis of extracellular miR-21 in A549 cells following treatment with the indicated PRIME-miR-21 formulations and free-component controls. For panels d and e, cells were treated with 100 nM PRIME at 37 °C for 48 h. RT–qPCR data are mean ± s.d. from three biological replicates.

Extracellular miRNA abundance was further quantified by RT–qPCR after 48 h of PRIME treatment. No exogenous miRNA was added in these assays. PRIME constructs against miR-21, miR-10, miR-23, miR-92 or miR-182 reduced their matched extracellular targets by 70.6–89.4% across A549, HeLa and MDA-MB-231 cells, whereas non-matched PRIME–miRNA combinations produced little depletion (Fig. 4d and Supplementary Fig. 11). To define the structural necessities for PRIME-mediated extracellular miRNA depletion, we next performed RT-qPCR analysis of extracellular miR-21 after treating A549 cells with different PRIME-miR-21 variants. Removal of either RNase H or the hairpin recognition unit markedly impaired miR-21 depletion, supporting the contribution of both components under the tested conditions (Fig. 4e and Supplementary Fig. 12). Constructs lacking the anchoring configuration were also less effective. The aptamer-directed formulation used a MUC1 aptamer as the targeting unit to recognize the cell-surface mucin MUC1, which has been reported for aptamer-mediated tumor-cell targeting in A549 and MDA-MB-231 models.^39,40^ MUC1 is also expressed and functionally targetable in H1299 NSCLC cells, supporting the use of MUC1 as a targeting marker in the orthotopic H1299 model.^41^ Thus, membrane-anchored PRIME depleted endogenous extracellular miRNAs in a sequence-selective manner over the 48 h treatment period.

### PRIME treatment reduces tumour-cell migration

We next tested whether depletion of extracellular miRNAs by membrane-anchored PRIME could suppress tumour-cell migration. Candidate miRNA targets were prioritized by integrating tumour-associated miRNA abundance from TCGA ^42^, supportive evidence of extracellular occurrence from GEO datasets ^43^ and reported associations with migration or metastasis^44,45,46^. This analysis identified miR-21, miR-10 and miR-23 as the principal combination for lung cancer, miR-21, miR-10 and miR-182 for breast cancer, and miR-21, miR-10 and miR-92 for cervical cancer (Supplementary Fig. 13).

The functional consequences of single-target depletion were examined in A549 cells using membrane-anchored PRIME-miR-21. PRIME inhibited migration by 76.6% in the wound-healing assay and by 40.3% in the Transwell assay (Fig. 5a and 5b). PRIME constructs lacking RNase H, the hairpin recognition unit or stable membrane anchoring agents showed substantially weaker activity. The differences in migration inhibition were consistent with the extracellular miR-21 depletion results (Fig. 4e). These findings suggest that PRIME relies on miRNA recognition, RNase H-mediated cleavage and stable membrane anchoring to suppress migration. Moreover, wound-healing and Transwell assays showed that extracellular miR-21 supplementation enhanced A549-cell migration. PRIME-miR-21 attenuated the migration increase associated with added miR-21 (Supplementary Fig. 14). CCK-8 and Annexin V/PI assays assessed metabolic activity and apoptosis, respectively. M-PRIME reduced the CCK-8 signal by 34.4%; limited apoptosis therefore does not exclude growth or metabolic effects on migration readouts (Supplementary Figs. 15 and 16). Changes in pro-migratory programmes, including epithelial–mesenchymal transition and cytoskeletal remodelling may also contribute ^44,45,46^.

**Figure 5.**
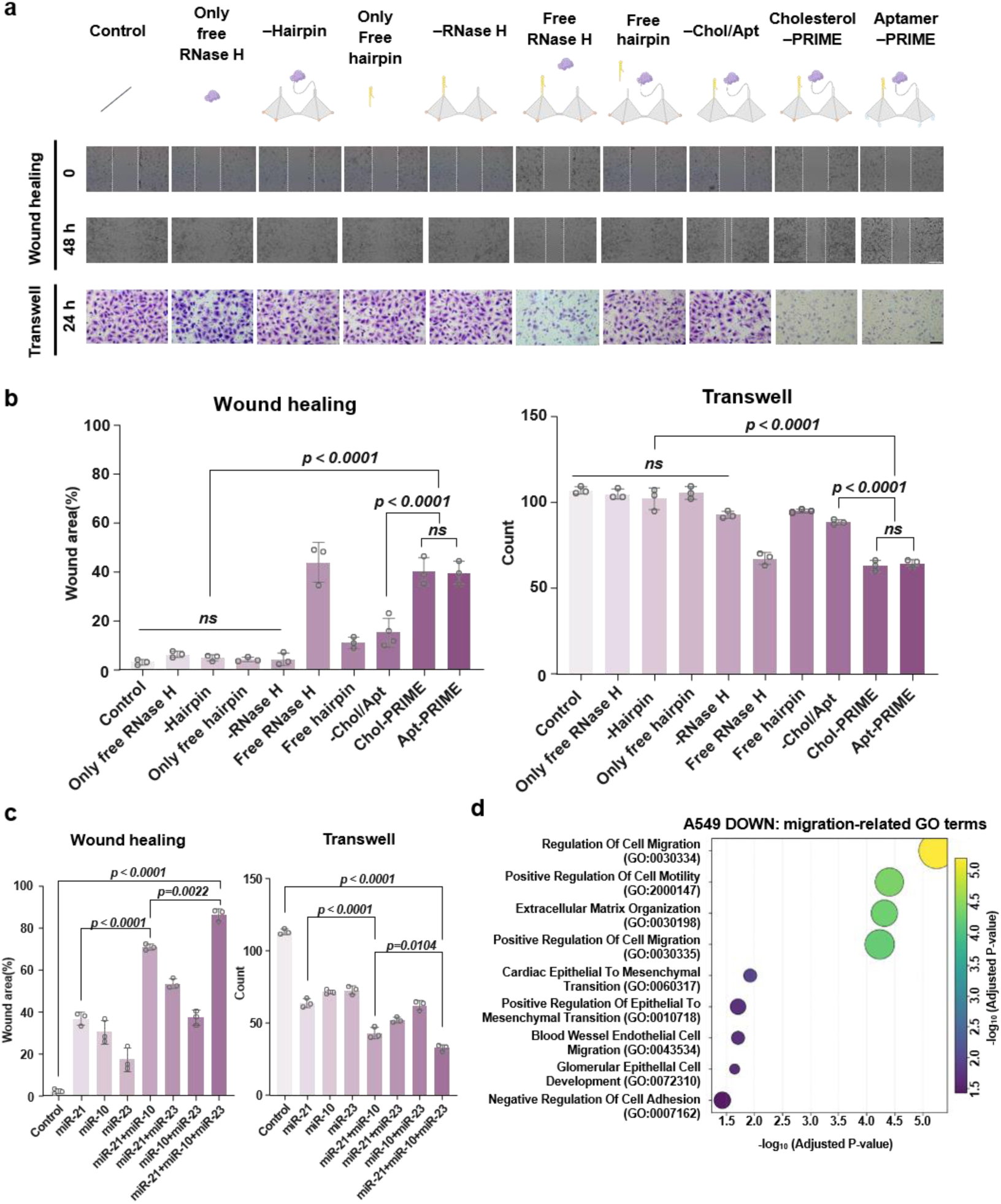
PRIME treatment reduces A549-cell migration. a, Representative wound-healing and Transwell images of A549 cells treated with the indicated PRIME-miR-21 formulations or free-component controls. b, Quantification of remaining wound area and migrated-cell numbers corresponding to the conditions in panel a. c, Quantification of A549-cell migration following treatment with individual PRIME constructs targeting miR-21, miR-10 or miR-23, dual-target combinations, or the three-target formulation M-PRIME comprising all three constructs. Wound-healing images were acquired at 0 and 48 h, and Transwell migration was assessed after 24 h. For corresponding images see Supplementary Fig. 17. d, Gene ontology enrichment analysis of genes downregulated in A549 cells after 48 h of treatment with M-PRIME targeting miR-21, miR-10 and miR-23. Enriched terms include biological processes associated with cell migration, epithelial–mesenchymal transition and extracellular-matrix organization. Bubble size indicates the number of downregulated genes overlapping each GO term, whereas bubble colour and the x axis indicate −log10(adjusted P value). Migration data are mean ± s.d; wound-healing assays and Transwell assays used three biological replicates. RNA sequencing was performed using three biological replicates per group. PRIME was used at 100 nM, and assays were performed at 37 °C. Wound-healing and RNA-seq endpoints were collected after 48 h; Transwell migration was assessed over 24 h with PRIME maintained throughout the migration period. Representative images are shown with scale bars of 100 μm for wound-healing assays and 20 μm for Transwell assays.

Because metastatic behaviour is regulated by interacting miRNA networks, we next tested whether the depletion of multiple prioritized extracellular targets could strengthen the response. All individual PRIME formulations against miR-21, miR-10 or miR-23 reduced A549-cell migration. Athree-target formulation comprising all three constructs (M-PRIME) produced the strongest effect (Fig. 5c and Supplementary Fig. 17). In the Transwell assay, migration inhibition increased from 40.3% with PRIME-miR-21 alone to 71.3% with M-PRIME. By contrast, a three-target control formulation directed against the low-ranked extracellular miRNAs miR-4438, miR-4665 and miR-3686 did not significantly inhibit migration (Supplementary Fig. 18).

RNA sequencing was used to examine the transcriptional changes associated with M-PRIME treatment. M-PRIME-treated A549 cells separated from untreated controls by principal-component analysis and displayed a reproducible differential-expression profile. Gene ontology analysis of downregulated genes identified coordinated changes in programmes associated with cell migration, epithelial–mesenchymal transition and extracellular-matrix organization (Fig. 5d and Supplementary Figs. 19–21). These transcriptomic data are consistent with broad remodelling of migration-associated cell states following extracellular miRNA depletion, but do not by themselves establish direct regulatory relationships between individual miRNAs and specific downstream genes.^4,5^

The functional effect of M-PRIME was subsequently tested across various types of tumour cell. M-PRIME directed against miR-21, miR-10 and miR-23 in NCI-H1299 lung cancer cells, miR-21, miR-10 and miR-182 in MDA-MB-231 breast cancer cells, or miR-21, miR-10 and miR-92 in HeLa cervical cancer cells consistently suppressed migration in both wound-healing and Transwell assays (Supplementary Fig. 22). Transwell migration was inhibited by 90.4% in H1299 cells, 96.0% in MDA-MB-231 cells and 87.0% in HeLa cells. These results show that PRIME can suppress migration across various tumour-cell models when programmed with the corresponding miRNA target combinations.

### Aptamer-directed M-PRIME enhances gemcitabine response in orthotopic lung tumours

To assess M-PRIME in established lung tumours, mice bearing orthotopic H1299-Luc tumours were randomly assigned to intravenous PBS, gemcitabine or combination-treatment groups. In the combination group, aptamer-directed M-PRIME was administered first, followed by gemcitabine 48 h later; this treatment cycle was repeated once weekly for four weeks. Treatment began approximately five weeks after intrapulmonary tumour-cell implantation (*n* = 3 mice per group; Fig. 6a). M-PRIME contained equal mass proportions of constructs targeting miR-21, miR-10 and miR-23; gemcitabine and M-PRIME were administered at 50 and 10 mg kg⁻¹, respectively. Longitudinal bioluminescence showed progressive pulmonary tumour expansion with PBS, partial restraint with gemcitabine, and a greater, sustained response to the gemcitabine/M-PRIME combination (Fig. 6b, 6c). Endpoint *ex vivo* lung imaging showed a lower pulmonary tumour burden with gemcitabine/M-PRIME than with gemcitabine alone (*p* = 0.0006; Fig. 6d). H&E-stained whole-lung sections showed numerous spatially distributed tumour nodules with PBS, a moderate reduction with gemcitabine, and the lowest burden after gemcitabine/M-PRIME combination therapy. Quantification of H&E-stained lung section confirmed fewer nodules with the combination therapy than with gemcitabine alone (*p* = 0.0113; Fig. 6e,f).

**Figure 6.**
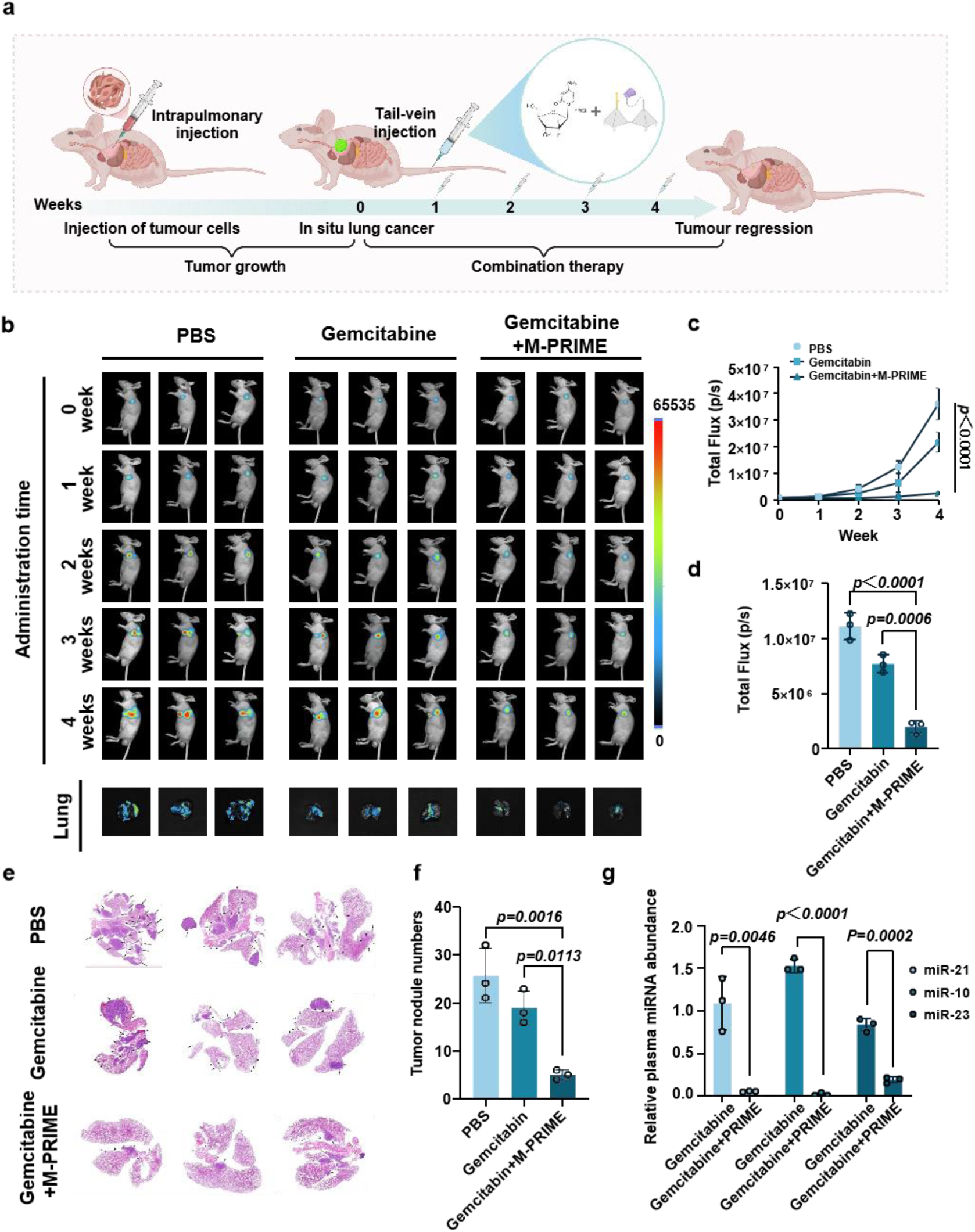
Aptamer-directed M-PRIME potentiates gemcitabine response in established orthotopic lung tumours. Equal mass proportions of aptamer-directed PRIME constructs targeting miR-21, miR-10 and miR-23 were combined, corresponding to approximately 3.33 mg kg^−1^ of each construct at a total M-PRIME dose of 10 mg kg^−1^. a, Orthotopic H1299-Luc lung-tumour treatment scheme. Approximately five weeks after intrapulmonary implantation, mice received intravenous PBS or gemcitabine (50 mg kg^−1^), whereas the combination group received M-PRIME (10 mg kg^−1^ total) first, followed by gemcitabine 48 h later; this treatment cycle was repeated once weekly for four weeks. b, Representative longitudinal whole-animal bioluminescence and endpoint *ex vivo* lung images. c, Pulmonary bioluminescence trajectories during treatment. d, Endpoint ex vivo lung bioluminescence as an index of pulmonary tumour burden. e, Representative H&E-stained whole-lung sections. The entire lung tissue area was used for tumour-nodule assessment, and arrows indicate representative tumour nodules. f, Tumour-nodule counts from one H&E-stained lung section per mouse. g, Plasma miR-21, miR-10 and miR-23 measured by RT–qPCR after gemcitabine or combination treatment. Values are mean ± s.d. from three mice per group. Bar graphs in panels d and f were analysed by one-way ANOVA with Tukey’s multiple-comparisons test; longitudinal bioluminescence was assessed by two-way repeated-measures ANOVA with Tukey’s multiple-comparisons test. All tests were two-sided; exact *p* values are reported in the panels. For panel g, each miRNA was compared between the gemcitabine and gemcitabine + M-PRIME groups using a two-sided unpaired Student’s t-test. Unadjusted *p* values are shown in the panel; *p* values were additionally adjusted across the three miRNA comparisons using the Holm–Šídák method.

Lower pulmonary tumour burden was accompanied by lower circulating target-miRNA levels. Combination treatment reduced circulating miR-21, miR-10 and miR-23 by 95.0%, 98.7% and 77.5%, respectively, relative to gemcitabine alone (Fig. 6g and Supplementary Fig. 23). Tumour sections also showed reduced EGR1, FOSL1, HAS3, CEMIP, TUBB3 and CDH11 signals, consistent with the migration- and matrix-associated expression changes observed *in vitro* (Supplementary Fig. 24). These endpoint measurements associate lower circulating miRNA levels with treatment response. The combination treatment was generally well tolerated, as assessed by body weight (Supplementary Fig. 25), haematology (Supplementary Fig. 26), serum biochemistry (Supplementary Fig. 27), plasma IgM, IgG, IL-6 and TNF-α levels (Supplementary Fig. 28), and major-organ histology (Supplementary Fig. 29). Within this proof-of-concept cohort, combining aptamer-directed M-PRIME with gemcitabine was associated with lower circulating target-miRNA levels and improved control of established pulmonary tumours.

### Early aptamer-directed M-PRIME restricts hematogenous breast-cancer lung colonization

The breast-cancer study addressed a different therapeutic window: interception before pulmonary colonies were established. In the MDA-MB-231-Luc tail-vein model, aptamer-directed M-PRIME targeting miR-21, miR-10 and miR-182 was administered one day before tumour-cell inoculation. In each weekly combination-treatment cycle, M-PRIME was administered first, followed by gemcitabine 48 h later; treatment was continued for four weeks.; PBS and gemcitabine groups followed corresponding schedules (Fig. 7a). Gemcitabine and M-PRIME were given at 50 and 10 mg kg⁻¹, respectively. This pre-inoculation design models post-intravasation survival and lung colonization, not regression of established metastases or dissemination from a primary mammary tumour.

**Figure 7.**
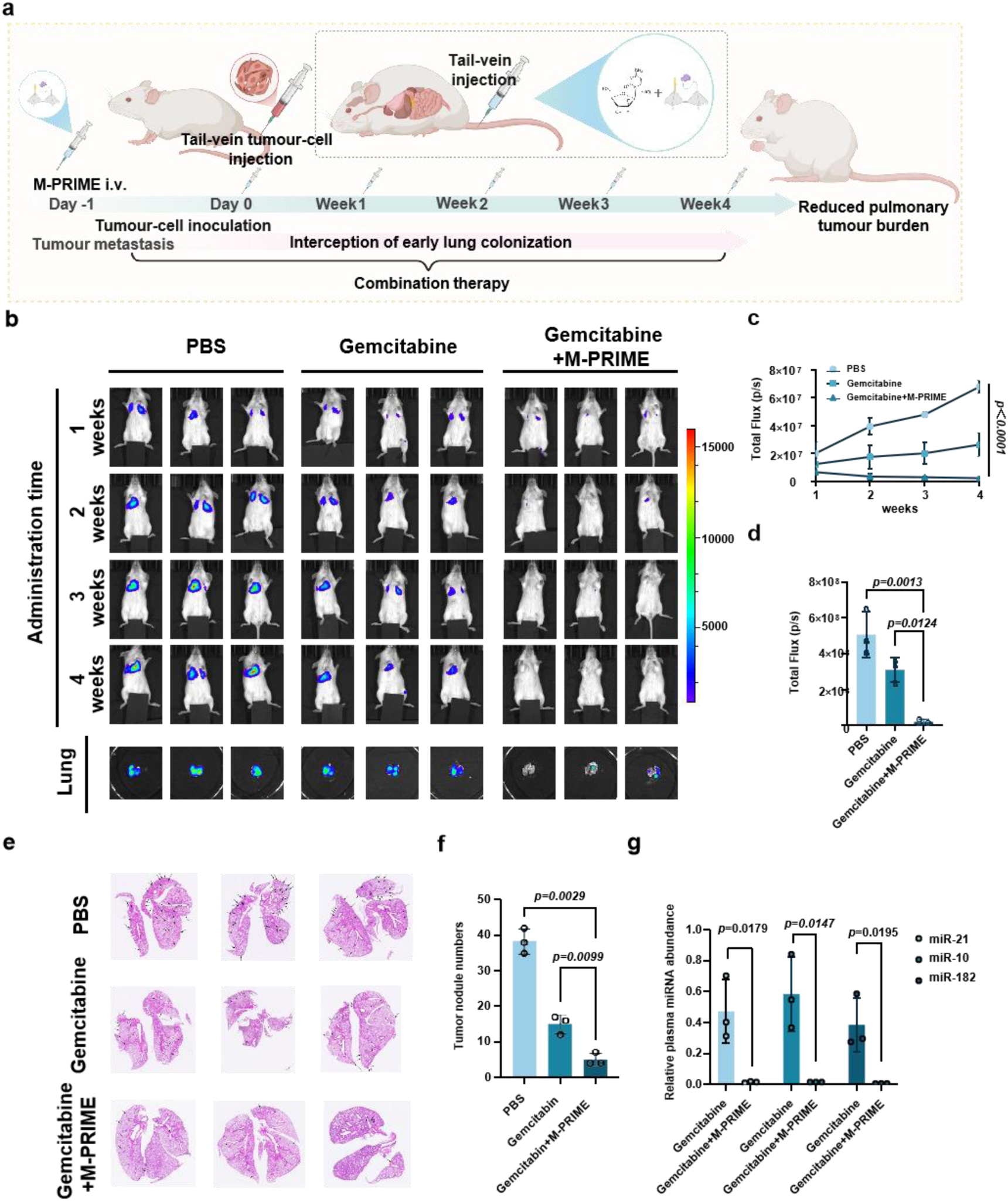
Early aptamer-directed M-PRIME restricts haematogenous breast-cancer lung colonization during gemcitabine treatment. M-PRIME contained equal proportions of constructs directed against miR-21, miR-10 and miR-182. a, Timing of intervention in the MDA-MB-231-Luc tail-vein model. M-PRIME (10 mg kg^−1^ total) was administered one day before tumour-cell inoculation. In each weekly combination-treatment cycle, M-PRIME was administered first, followed by gemcitabine (50 mg kg^−1^) 48 h later, for four consecutive weeks.; PBS and gemcitabine controls followed matched schedules. This design assesses early lung colonization after intervention before tumour-cell inoculation. b, Serial in vivo bioluminescence images, followed by endpoint imaging of excised lungs. c, Four-week changes in pulmonary bioluminescence. d, Final ex vivo lung signal as an index of metastatic burden. e, Whole-lung H&E sections illustrating colonization across treatment groups. The entire lung tissue area was used for tumour-nodule assessment, and arrows indicate representative tumour nodules. f, Pulmonary tumour-nodule counts from one H&E-stained lung section per mouse. g, Circulating miR-21, miR-10 and miR-182 quantified by RT–qPCR after gemcitabine or combination treatment. Results are mean ± s.d. (n = 3 mice per group). One-way ANOVA with Tukey’s multiple-comparisons test was applied to bar graphs in panels d and f, whereas longitudinal bioluminescence was evaluated by two-way repeated-measures ANOVA with Tukey’s multiple-comparisons test. Tests were two-sided, with exact *p* values displayed in the relevant panels. For panel g, each miRNA was compared between the gemcitabine and gemcitabine + M-PRIME groups using a two-sided unpaired Student’s t-test. Unadjusted *p* values are shown in the panel; *p* values were additionally adjusted across the three miRNA comparisons using the Holm–Šídák method.

Early M-PRIME markedly altered the subsequent colonization trajectory. Lung bioluminescence remained lower with the combination than with gemcitabine alone throughout follow-up (*p* < 0.0001; Fig. 7b, 7c). Endpoint imaging showed 95.8% inhibition relative to PBS, compared with 38.7% for gemcitabine, and confirmed a lower pulmonary burden in the combination group (*p* = 0.0124; Fig. 7d). Histology corroborated this effect: tumour-nodule inhibition reached 87.0% with the combination versus 60.9% with gemcitabine, with fewer nodules in the combination group (*p* = 0.0099; Fig. 7e, 7f). Thus, intervention before tumour-cell challenge restricted the establishment and expansion of intravenously disseminated breast-cancer cells in the lung.

Circulating miRNA levels decreased alongside the reduction in lung colonization. Combination treatment reduced circulating miR-21, miR-10 and miR-182 by 96.6%, 97.2% and 98.2%, respectively, relative to gemcitabine alone (Fig. 7g and Supplementary Fig. 30). The combination treatment was generally well tolerated in this model, as assessed by body weight (Supplementary Fig. 31), haematological parameters (Supplementary Fig. 32), liver and kidney function indices (Supplementary Fig. 33), and plasma IgM, IgG, IL-6 and TNF-α levels (Supplementary Fig. 34). Together with the orthotopic lung-tumour study, these data show that aptamer-directed M-PRIME can strengthen gemcitabine response at distinct disease stages—against established pulmonary disease or during early lung colonization.

## Discussion

PRIME enables sustained depletion of extracellular miRNAs by organizing target recognition and enzymatic cleavage at the cell surface. The double-tetrahedral scaffold co-localizes an exchangeable recognition hairpin and RNase H and supports membrane anchoring. Separate assays showed repeated target-induced reaction cycles, membrane-associated signals persisting for 48 h and depletion of matched extracellular miRNAs by 70.6–89.4% across three cancer-cell models. M-PRIME treatment was associated with the inhibition of cell Transwell migration and downregulation of migration- and matrix-associated programmes. *In vivo*, aptamer-directed M-PRIME reduced circulating targets by 77.5–98.7% and, when combined with gemcitabine, lowered pulmonary tumour burden by 74.5% in established orthotopic lung tumours and by 93.2% during early breast-cancer lung colonization relative to gemcitabine alone. The central advance is the use of a membrane-anchored DNA scaffold to sustain sequence-selective extracellular RNA degradation.

PRIME differs from most RNA-targeting technologies in acting on extracellular miRNAs at the cell surface. Antagomirs, locked-nucleic-acid or antisense inhibitors and miRNA sponges are readily programmable, but generally require cellular uptake and suppress miRNAs through stoichiometric binding or intracellular RNase H recruitment^15,16,17,18^. Cas13 systems and emerging RiboTACs introduce catalytic or induced RNA degradation, yet operate predominantly inside cells, requiring delivery or expression of a nuclease–guide complex or recruitment of an endogenous ribonuclease, with continued concerns about delivery efficiency, collateral cleavage and intracellular off-target effects^19,20,47,48^. At the extracellular level, recent exosome-disruption strategies intercept pathological communication but alter whole vesicle populations, whereas membrane-anchored DNA devices have mainly been developed for surface sensing, logic computation and triggered cargo release^49,50^. PRIME uses this membrane-associated format to couple programmable RNA recognition to RNase H-mediated cleavage, with repeated reporter responses and lower extracellular target-miRNA levels measured in separate assays. The design addresses accessible extracellular miRNAs. Further development will require reproducible manufacture of the enzyme–DNA conjugate and characterization of its *in vivo* catalytic lifetime, biodistribution and efficacy as a monotherapy.

The most immediate opportunity is to develop PRIME from fixed multi-miRNA formulations into disease- and patient-matched extracellular-RNA depletion systems. Recognition panels could be selected from longitudinal plasma and extracellular-vesicle RNA profiles and exchanged without redesigning the scaffold, allowing target combinations to follow tumour type, treatment pressure and disease stage^7,14,51,52^. In parallel, improving nuclease resistance, optimizing enzyme loading and targeting-ligand density, and defining dose–membrane occupancy–RNA-turnover relationships should increase durability and manufacturability. These advances should be evaluated in larger cohorts that include M-PRIME monotherapy, immunocompetent hosts, spontaneous metastasis and treatment of established metastatic lesions. With these advances, sequence-selective extracellular RNA degradation could also be explored in inflammatory, fibrotic and other disorders in which extracellular RNAs contribute to pathological cell–cell communication^53,54,55,56^.

## Methods

### Reagents and materials

Oligonucleotides used for PRIME scaffold assembly, target recognition and fluorescence reporting are listed in Supplementary Table 1. DNA and RNA oligonucleotides were synthesized by Sangon Biotech (Shanghai, China) and purified by PAGE, ULTRAPAGE or HPLC as appropriate for the modification and application. RNase H (BestEnzymes, cat. no. EG20208S; 50 U μl⁻¹) was used with 1× RNase H Reaction Buffer (New England Biolabs), containing 50 mM Tris-HCl, 75 mM KCl, 3 mM MgCl₂ and 10 mM DTT (pH 8.3 at 25 °C). DNase/RNase-free water, nuclease-free tubes and filtered tips were used for nucleic-acid handling. Commercial chemicals were used as received unless a purification step is specified below.

### Oligonucleotides and cell lines

Sequences, chemical modifications, and functional roles of oligonucleotides used for PRIME construction and fluorescence assays are provided in Supplementary Table 1. Stock solutions were prepared in nuclease-free water, aliquoted and stored to avoid repeated freeze-thaw cycles. A549, H1299, HeLa and MDA-MB-231 cells were cultured in RPMI-1640 medium supplemented with 10% fetal bovine serum and 1% penicillin-streptomycin at 37 °C in a humidified incubator containing 5% CO_2_. Cells were passaged at approximately 80-90% confluence using 0.25% trypsin-EDTA. Luciferase-expressing H1299-Luc and MDA-MB-231-Luc cells were maintained under the same conditions before animal-model construction.

### Preparation of RNase H-DNA conjugates

RNase H was covalently linked to thiol-modified DNA using Sulfo-SMCC chemistry. RNase H in PBS (pH 7.2–7.4) was reacted with a 10-fold molar excess of Sulfo-SMCC for 1 h at room temperature with gentle agitation to introduce maleimide groups on the protein. Unreacted Sulfo-SMCC and other low-molecular-weight components were removed using a Zeba Spin Desalting Column (7K MWCO; Thermo Fisher Scientific). The maleimide-activated RNase H was then mixed with pre-reduced thiol-modified SH-DNA at a DNA:RNase H molar ratio of 1:5 and reacted at 4 °C for 8 h to form the RNase H–DNA conjugate. The reaction mixture was subsequently purified using an Amicon Ultra-0.5 centrifugal filter (30 kDa MWCO; Merck Millipore), with the retained conjugate recovered by reverse centrifugation, to reduce unconjugated DNA, free RNase H and low-molecular-weight components. The purified conjugate was aliquoted and stored at −20 °C until use. RNase H, SH-DNA and RNase H–DNA conjugate samples were mixed with matrix solution, spotted onto the MALDI target plate and analyzed in positive-ion mode. Spectra were acquired under the same instrument settings for the unconjugated components and the conjugate. Observed mass peaks were used to assess RNase H– DNA conjugate formation.

### Preparation and characterization of PRIME

Hairpin recognition units were designed using NUPACK^57^ to generate hairpins with a 10-bp stem, a 12-nt loop and an 8-nt toehold segment. The target-complementary region was matched to the cognate miRNA and designed to open upon target hybridization while maintaining a stable closed hairpin at 37 °C. Candidate hairpins were screened to reduce undesired secondary structures and to maintain favorable free-energy separation between the closed and target-bound states.

PRIME was assembled from two DNA tetrahedral subunits. The recognition tetrahedral subunit was assembled from S1, S2-HP, S3 and S4, after which the target-specific hairpin (HP) recognition strand was hybridized to the complementary region of S1, whereas the catalytic tetrahedral subunit was assembled from S1, S2-SH, S3 and S4. Each subunit was assembled by mixing the corresponding oligonucleotides at an equimolar ratio in TM buffer (10 mM Tris-HCl, 50 mM MgCl₂, pH 8.0), heating at 85 °C for 10 min and cooling to 4 °C for 30 min. The two preassembled subunits were then mixed at an equimolar ratio and incubated at 37 °C for 1 h to hybridize the complementary linker regions, yielding the double-tetrahedral PRIME scaffold. The RNase H-DNA conjugate was subsequently incubated with the assembled tetrahedral scaffold at 37 °C for 1 h to complete catalytic-unit loading and final PRIME assembly. Cholesterol-modified strands were incorporated for cell-surface anchoring, and aptamer-bearing strands were incorporated for in vivo tumor targeting.

Scaffold assembly, RNase H-DNA conjugate loading and serum stability were characterized by native PAGE. For scaffold-assembly verification, samples were prepared at 1 μM, diluted to 100 nM, mixed with 6× loading buffer at a 5:1 volume ratio and loaded onto 4%–20% gradient native polyacrylamide gels. Electrophoresis was performed in an ice-cooled chamber at 90 V for 60 min unless otherwise stated, followed by SYBR Gold staining and UV gel imaging.

For AFM characterization, freshly cleaved mica was used as the substrate. DNA scaffold or assembled PRIME samples were diluted with ultrapure water, deposited onto mica, incubated at room temperature to allow adsorption, gently rinsed to remove salts and unbound species, and dried under nitrogen or air. Samples were imaged in tapping mode using a Bruker Dimension Edge atomic force microscope. Height images were used for morphology inspection and Feret-diameter analysis of well-isolated particles.

### RNase H activity assay

RNase H substrate selectivity was evaluated using various nucleic-acid substrates. Each substrate was incubated either in the absence or presence of RNase H under identical reaction conditions. Changes in substrate bands following RNase H treatment were assessed by native PAGE. Electrophoresis was performed on a 10% native PAGE gel at 120 V for 40 min.

### Coarse-grained oxDNA simulations

Coarse-grained structural models were built in oxView and simulated using oxDNA^35,36^. The double-tetrahedral scaffold, hairpin recognition unit, linker domains and SH-DNA arm were included in the input model. Topology (.top) and configuration (.dat) files were exported from oxView. Simulations were performed at 37 °C in a 100 × 100 × 100 simulation box (1 length unit = 0.8518 nm). Each trajectory was propagated for 500,000 molecular-dynamics steps using an integration time step of 0.001 simulation units. Defined nucleotide positions were used to calculate Euclidean distances across the trajectory. A-B, C-D and E-F distances were used to estimate global scaffold dimensions and linker geometry, whereas G1 and G2 defined the two ends of the miRNA-bound recognition region and H was defined as nucleotide 60 of SH-DNA, corresponding to the 3′-terminal nucleotide adjacent to the C6 thiol linker used for RNase H conjugation. Root-mean-square fluctuation analysis was used to evaluate local rigidity and flexibility across scaffold regions. RNase H itself and the Sulfo-SMCC/C6 linker were not explicitly represented in the oxDNA model; consequently, no protein-derived steric, orientational or conformational constraints were imposed at H. H therefore represents only the scaffold attachment position and does not model the structure or catalytic orientation of RNase H.

### Bioinformatic prioritization of extracellular pro-metastatic miRNAs

Candidate extracellular pro-metastatic miRNAs were prioritized by integrating tumour-associated miRNA abundance, extracellular-miRNA evidence and literature-supported migration or metastasis annotations. Mature-miRNA expression matrices from the TCGA LUAD, BRCA and CESC cohorts were processed using miRBase nomenclature^58^. Sample-level miRNA counts were normalized as reads per million mapped mature-miRNA reads (RPM). TCGA samples with barcode suffixes −01 or −02 were classified as tumour samples, whereas −10 or −11 samples were classified as normal samples for tumour–normal comparisons. Under this classification, the processed datasets comprised 450 tumour and 45 normal samples for LUAD, 749 tumour and 76 normal samples for BRCA, and 306 tumour and 3 normal samples for CESC. Seven BRCA samples and two CESC samples with barcode suffix −06 were not assigned to either the tumour or normal group for tumour–normal comparisons. No fixed low-expression cutoff was applied; instead, the 50 most abundant miRNAs ranked by mean expression within each processed TCGA cohort were retained as the abundance-prefiltered set. These miRNAs were intersected with disease-specific literature annotations of migration or metastasis, retaining miRNAs annotated as positively associated with these phenotypes. Extracellular-miRNA evidence was obtained from 36 NSCLC plasma profiles from 18 patients in GSE244643, all 12 breast-cancer plasma samples in GSE240872 and 24 plasma samples in GSE302097. miRNA names were harmonized across datasets before intersection. Candidates supported by TCGA abundance, extracellular occurrence and migration- or metastasis-associated annotations were retained and ranked by mean TCGA RPM within each cancer context to define the final PRIME target sets.

### Fluorescence kinetic assays of PRIME-miRNA reactions

Fluorophore-quencher-labeled PRIME constructs were used for solution-phase fluorescence assays. Cy5 fluorescence was quenched by BHQ3 in the closed hairpin state and recovered after target-induced hairpin opening. Reactions were performed in 20 μl volumes at 37 °C using 100 nM PRIME and 100 nM target miRNA on a real-time fluorescence qPCR instrument with red-channel detection and a 5 s acquisition cycle. For standard measurements, target miRNA was added immediately before rapid centrifugation and fluorescence acquisition. For repeated-input experiments, acquisition was paused at 10-min intervals and 1 μl of target-miRNA solution was added at each time point. Each input was formulated so that the newly added target miRNA reached a final concentration of 100 nM in the updated total reaction volume immediately after addition. Thus, 100 nM refers to the final concentration of each newly introduced miRNA input rather than the cumulative target concentration. The total reaction volume was updated after each 1 μl addition, and concentrations were calculated using the post-addition volume to account for cumulative dilution. Acquisition was resumed immediately after rapid centrifugation.

For specificity analysis, five exchangeable-unit PRIME variants (PRIME-miR-21, PRIME-miR-10, PRIME-miR-23, PRIME-miR-182 and PRIME-miR-92) were tested with their cognate miRNAs and a panel of non-cognate miRNAs, with each PRIME variant and miRNA input used at 100 nM. Fluorescence–time curves were recorded, and the area under the curve (AUC) from each individual PRIME/miRNA reaction was used as an integrated response metric. For heat-map visualization, AUC values were normalized to the cognate-target response for each PRIME variant. Fluorescence data were processed in Excel and visualized using GraphPad Prism.

### Fluorescence microscopy for characterization of cell membrane functionalization of PRIME

Cells were seeded in confocal dishes at 5 × 10^4^ cells ml^−1^ in 2 ml medium and cultured for 1–2 d before treatment. When cell density reached approximately 30% or higher, cells were washed three times with PBS and incubated with 300 nM PRIME, 1× Hoechst 33342 and 1× WGA in cell-culture medium or buffer for 30 min at 37 °C. After incubation, cells were washed three times with PBS to remove unbound PRIME and excess dyes, and D-PBS was added for imaging.

Single-time-point fluorescence images were acquired after bright-field focusing in the same field of view for nuclear, membrane and PRIME fluorescence channels. For live-cell time-lapse imaging, dishes were placed in an environmental imaging chamber. Membrane and nuclear images were first acquired, and the Cy5 channel was then imaged at 10 min intervals. The excitation shutter was closed between acquisitions to reduce photobleaching and phototoxicity. Target miRNA was added to a final concentration of 100 nM before the first PRIME-channel image, and a second miRNA input was added after 100 min in repeated-input experiments. Cy5-labeled unquenched cholesterol-modified PRIME was used for membrane loading and retention assays, whereas fluorophore–quencher-labeled PRIME was used for target-response imaging. Membrane-associated fluorescence was quantified in ImageJ. Bright-field images were used to define cell boundaries, and a membrane-associated region of interest with a 2-pixel width was generated around each cell boundary. The same region of interest was applied to the corresponding fluorescence channel. Background subtraction and normalization were applied consistently across groups.

### Flow cytometry

For flow-cytometric analysis of membrane loading and cell-surface fluorescence responses, cells were grown to approximately 90% confluence, washed with PBS and detached with 0.25% trypsin-EDTA. Digestion was stopped with serum-containing medium, and cells were collected by centrifugation at 1,000 rpm for 3 min. Cells were resuspended in PBS or D-PBS and adjusted to 1 × 10^6^ cells ml^−1^. Cells were incubated with 300 nM PRIME for 30 min at 37 °C, centrifuged, resuspended in 500 μl D-PBS and passed through a 40 μm cell strainer. FSC-A, SSC-A, FSC-H, SSC-H and APC/Cy5-channel signals were collected in low-flow mode, with 10,000 events recorded per sample. For target-response assays, 100 nM target miRNA was added to PRIME-functionalized cell suspensions, and samples were measured every 10 min for up to 100 min.

### Extraction and RT-qPCR quantification of extracellular miRNAs

Extracellular miRNAs from cell-culture supernatants and plasma were extracted using the TIANGEN miRcute serum/plasma miRNA extraction and isolation kit. Cell-culture supernatants were collected into RNase-free tubes, centrifuged at 1,000 rpm for 5 min to remove cells and debris, and 200 μl of each sample was used for extraction. cel-miR-39 spike-in control was added before chloroform extraction and column purification. miRNAs were eluted with 20 μl RNase-free water and stored at −80 °C until analysis. Purified miRNAs were reverse-transcribed using a tailing-based miRNA reverse-transcription kit at 37 °C for 60 min followed by 85 °C for 5 min. RT-qPCR was performed using SYBR Green qPCR Master Mix, target-specific forward primers and the kit-provided universal reverse primer for 50 cycles, followed by melting-curve analysis. Only reactions with a single melting peak were included. Relative extracellular miRNA abundance was calculated by the ΔΔCt method using cel-miR-39 as the external reference^59^. Cell-culture depletion assays used three biological replicates. For cell-culture depletion assays, A549, HeLa or MDA-MB-231 cells were seeded in six-well plates, washed three times with PBS at approximately 60–70% confluence and treated for 48 h with complete medium containing a total PRIME concentration of 100 nM. PRIME variants were tested in the corresponding cell models as indicated in the figure legends. For component-control experiments, the nominal molar amounts of catalytic and recognition units were matched to those present in the complete PRIME treatment. Thus, for comparison with 100 nM single-target PRIME, free RNase H and free recognition hairpin were each added at an equivalent molar concentration of 100 nM, and unanchored PRIME was used at 100 nM. Cells were washed three times with PBS before treatment. No additional wash or medium replacement was performed after addition of the soluble components, so that free RNase H, free hairpin and unanchored PRIME remained in the medium throughout the 48-h incubation. Matching was based on one RNase H catalytic unit and one target-recognition hairpin per PRIME construct.

### Wound-healing migration assays

Cells were serum-starved before migration assays to reduce the influence of serum-derived growth factors. For wound-healing assays, cells were cultured in serum-free medium for 24 h before seeding or treatment. Cells were seeded in six-well plates with premarked reference lines and allowed to adhere until a confluent monolayer formed. The monolayer was washed with PBS, and a straight scratch was generated using a sterile 20–200 μl pipette tip. Detached cells were removed by washing twice with PBS, and 2 ml of treatment medium containing the indicated PRIME formulation at 100 nM was added. For exogenous miR-21 supplementation experiments, synthetic miR-21 was added to the wound-healing culture medium at a final concentration of 100 nM in 2 ml per well (200 pmol per well); in the corresponding Transwell experiments, miR-21 was added at 100 nM to both the 200 μl upper chamber and the 500 μl lower chamber. Images were acquired immediately after scratch generation (0 h) and after 48 h at four fixed positions per well using a 5× objective. Wound areas were measured using ImageJ, and remaining wound area (%) was calculated as Aₜ/A₀ × 100%, where A₀ and Aₜ represent the wound areas at 0 h and the indicated time point, respectively.

### Transwell migration assays

For Transwell assays, cells were serum-starved for 24 h before the migration assay, and a preliminary optimization was performed for each cell line to select cell seeding density and migration time. Serum-starved cells were resuspended in serum-free medium at 2 × 10^5^ to 6 × 10^5^ cells ml^−1^, and 200 μl cell suspension was added to the upper chamber. The lower chamber contained 500 μl complete medium as a chemoattractant. Based on the preliminary experiments, 4 × 10^5^ cells ml^−1^ and 24 h migration were selected for the formal experiments. For PRIME-treated groups, the indicated PRIME formulations were added at 100 nM at the start of the Transwell assay and maintained throughout the 24 h migration period. After migration, non-migrated cells on the upper surface of the membrane were gently removed with a cotton swab. The inserts were then fixed in 4% tissue fixative for 30 min and stained with 10% crystal violet working solution for 10 min. Inserts were washed with ultrapure water, mounted on glass slides and imaged using an inverted microscope with a 20× objective. Three fields were captured per insert, and migrated cells on the lower membrane surface were segmented and counted using ImageJ. Migration inhibition rates were calculated from migrated-cell counts relative to the corresponding control group. Data were processed using GraphPad Prism.

### CCK-8 metabolic-activity assay

CCK-8 assays were used to evaluate changes in cellular metabolic activity after PRIME treatment. A549 cells were seeded in 96-well plates at 5 × 10^4^ cells ml^−1^, 100 μl per well, and allowed to adhere. Cells were washed twice with PBS and treated with medium containing the indicated PRIME formulations at a final concentration of 100 nM for 48 h. Treatment groups included untreated control, L-PRIME, single-target PRIME and M-PRIME. After treatment, 10 μl CCK-8 reagent was added to each well and incubated for 1.5 h at 37 °C in the dark. Absorbance was measured at 450 nm using a microplate reader. CCK-8 signal inhibition was calculated as [(ODcontrol − ODtreated)/ODcontrol] × 100% after blank subtraction.

### Apoptosis analysis

Apoptosis was assessed by Annexin V/PI staining. A549 cells were seeded in six-well plates, treated with the indicated PRIME formulations at 100 nM for 48 h and collected together with the corresponding culture supernatants to include detached cells. Adherent cells were detached using EDTA-free trypsin, neutralized with complete medium and combined with the supernatant. Cells were collected by centrifugation, washed with cold PBS and resuspended in 1× Annexin V binding buffer at approximately 1 × 10^6^ cells ml^−1^. For each sample, 100 μl cell suspension was incubated with 5 μl Annexin V-FITC for 15 min at room temperature in the dark, followed by addition of 5 μl PI immediately before acquisition. A total of 10,000 events were collected per sample. Debris was excluded by FSC/SSC gating, and viable, early apoptotic, late apoptotic or secondary necrotic, and PI-positive dead-cell populations were assigned according to Annexin V-FITC and PI staining. Data were analyzed using FlowJo.

### RNA sequencing and transcriptomic analysis

For RNA-seq, serum-starved A549 cells were treated for 48 h with 100 nM cholesterol-modified M-PRIME targeting miR-21, miR-10 and miR-23 or with control medium. Three biological replicates were prepared for each group. After treatment, cells were washed three times with PBS, collected by gentle scraping, pelleted by centrifugation at 2,000 rpm for 5 min and snap-frozen in liquid nitrogen or stored at −80 °C before RNA extraction. Total RNA was extracted and treated with DNase I to remove residual DNA. Poly(A)+ mRNA was enriched using oligo(dT) magnetic beads, fragmented to approximately 200–300 nt, and used for strand-specific library preparation with an MGI-compatible library-preparation kit (Vazyme, cat. no. NR605). Libraries were sequenced on an MGI T7 platform to generate paired-end reads. Raw paired-end reads in FASTQ format were processed using fastp v0.20.1. Reads containing sequencing adapters, reads with >5% ambiguous bases (N), and reads in which >20% of bases had quality scores below Q10 were removed. Clean reads were aligned to the human reference genome GRCh38 using HISAT2 v2.2.1 with the parameters --phred33 --sensitive --no-discordant --no-mixed - I 1 -X 1000. Gene-expression levels were quantified using StringTie v2.1.5 with the -e -B parameters and ballgown v2.18.0. Differential-expression analysis was performed using DESeq2 v1.26.0^60^, with treatment condition as the sole design factor (design = ∼ condition), to compare M-PRIME-treated samples with controls. Multiple-testing correction was performed using the Benjamini–Hochberg procedure. Genes with |log2(fold change)| ≥ 0.58 and adjusted P < 0.05 were retained as differentially expressed genes for downstream enrichment and heat-map analyses. For visualization in the volcano plot, genes with |log2(fold change)| ≥ 1 and adjusted P < 0.05 were highlighted as strongly up- or downregulated. Principal component analysis was used to evaluate overall sample separation and replicate consistency. Heat maps were generated from log2(FPKM + 1)-transformed expression values after row-wise Z-score transformation and hierarchical clustering using Euclidean distance and average linkage. Gene Ontology enrichment analysis was performed on downregulated differentially expressed genes (log2(fold change) ≤ −0.58 and adjusted P < 0.05) using GSEApy Enrichr with the GO_Biological_Process_2023 gene-set library^61^. No custom background gene set was specified, and the default Enrichr background was used. P values were adjusted for multiple testing using the Benjamini–Hochberg procedure, and terms with adjusted *p* < 0.05 were considered significantly enriched. Migration-, motility-, adhesion-, cytoskeleton-, extracellular-matrix-, epithelial–mesenchymal transition- and invasion-related biological processes were selected for visualization and migration-associated gene-set construction.

### Animal models and *in vivo* treatment

All animal experiments were performed in accordance with institutional animal-care guidelines and were approved by Ethics Committee of Beijing University of Chemical Technology (case number: BUCT21-2512-054). BALB/c-nu nude mice were used for the orthotopic H1299-Luc lung-tumor model, and NCG mice were used for the MDA-MB-231-Luc hematogenous metastasis-interception model. Mice were housed under specific-pathogen-free conditions at 22-26 °C and 40-60% relative humidity with a 12 h light/dark cycle and free access to food and water. Mice were acclimated for 1 week before experiments and randomly assigned to treatment groups. The in vivo cohorts were designed as proof-of-concept cohorts, with n = 3 mice per group unless otherwise stated.

H1299-Luc cells were collected during logarithmic growth, prepared as a single-cell suspension and resuspended in serum-free DMEM. Before inoculation, the cell suspension was mixed with Matrigel (Meiluncellgel, PWL306) at a 1:1 volume ratio to limit cell dispersion within the thoracic cavity and support localized orthotopic tumour establishment. Cell viability was checked by trypan blue exclusion before inoculation. Six- to seven-week-old male BALB/c-nu mice weighing 20–25 g were anesthetized with 1.25% Avertin, the thoracic region was disinfected, and 40 μl of the H1299-Luc/Matrigel suspension, corresponding to 8 × 10⁵ cells per mouse, was injected into the lung through the left axillary intercostal space using an insulin syringe. Mice were monitored during recovery and returned to the animal facility after normal breathing and activity were confirmed. Tumors were allowed to establish for approximately 5 weeks and were monitored weekly by in vivo bioluminescence imaging before treatment initiation.

For the breast-cancer haematogenous metastasis-interception model, 6–7-week-old female NCG mice weighing 15–17 g were used. MDA-MB-231-Luc cells were collected during logarithmic growth after confirmation of negative mycoplasma status, washed, resuspended in sterile PBS at 6 × 10⁶ cells ml⁻¹ and injected through the tail vein at 100 μl per mouse, corresponding to 6 × 10⁵ cells per mouse. NCG mice received aptamer-directed M-PRIME one day before tumor-cell inoculation. MDA-MB-231-Luc cells were then injected through the tail vein to initiate hematogenous dissemination. Initial bioluminescence imaging was performed 15 min after tumour-cell inoculation to record the initial distribution of the injected luciferase-expressing cells. Because the material was administered before tumor-cell inoculation, this model was interpreted as early metastasis interception rather than treatment of established breast metastases.

For *in vivo* treatment, mice received intravenous PBS, gemcitabine or gemcitabine plus aptamer-directed M-PRIME for four consecutive weeks. Gemcitabine was prepared in physiological saline at 10 mg ml^−1^ and administered at 50 mg kg^−1^ with the injection volume adjusted according to body weight. Aptamer-directed M-PRIME was prepared at a total formulation concentration of 2 mg ml^−1^ and administered at a total dose of 10 mg kg^−1^, calculated on the basis of the combined mass of the three complete target-specific PRIME constructs, with the injection volume adjusted according to body weight. In combination groups, M-PRIME was administered first, followed by gemcitabine 48 h later. This treatment cycle was repeated once weekly for four consecutive weeks. In vivo imaging was performed 72 h after gemcitabine administration in each weekly cycle, and blood and tissue samples were collected at the experimental endpoint. Lung-cancer M-PRIME contained aptamer-directed PRIME variants targeting miR-21, miR-10 and miR-23; breast-cancer M-PRIME contained aptamer-directed PRIME variants targeting miR-21, miR-10 and miR-182. The three PRIME variants in each formulation were mixed at equal mass proportions, corresponding to approximately 3.33 mg kg^−1^ of each target-specific PRIME construct at a total M-PRIME dose of 10 mg kg^−1^.

### *In vivo* and *ex vivo* bioluminescence imaging

Tumour-inhibition rates relative to PBS were calculated as [1 − (mean tumour burden of the treated group / mean tumour burden of the PBS group)] × 100%. Reductions in tumour burden relative to gemcitabine alone were calculated as [1 − (mean tumour burden of the combination group / mean tumour burden of the gemcitabine group)] × 100%. Tumor burden was monitored by bioluminescence imaging of luciferase-expressing tumor cells. D-luciferin was dissolved in sterile PBS and injected intraperitoneally at 150 mg kg^−1^. After approximately 10 min to allow substrate distribution, mice were anesthetized using isoflurane (2–3% for induction and 1.5–2% during imaging) and placed in the imaging chamber in a consistent position. Exposure time, binning and gain were kept constant within each experiment. Total flux (photons s^−1^; reported as “Total Flux [p/s]” in Living Image software) was quantified using circular regions of interest with consistent analysis settings. At the endpoint, lungs were harvested and imaged ex vivo using the same acquisition and analysis settings.

### Histology, pulmonary tumour-nodule counting and immunofluorescence staining

At the experimental endpoint, mice were anesthetized and euthanized according to approved procedures. Blood was collected, and major organs were harvested immediately. Tissues for histology were fixed in 4% tissue fixative for 24 h, dehydrated, paraffin embedded and sectioned at approximately 4 μm thickness. Sections were deparaffinized, rehydrated and stained with hematoxylin and eosin. Major organs including liver, spleen, lung, brain and kidney were examined for histopathological changes. One H&E-stained lung section per mouse was analyzed using ImageViewer, and pulmonary tumour nodules were counted across the entire lung tissue area of the section. Because only one section per mouse was used for nodule quantification, section-spacing criteria and cross-section deduplication of the same nodule were not applicable. Pulmonary tumour-nodule counting was performed by an investigator blinded to treatment-group identity.

For immunofluorescence staining of tumor or lung sections, paraffin sections were deparaffinized, rehydrated and subjected to EDTA-buffer antigen retrieval under heating. After cooling and PBS washing, sections were blocked with 5% BSA for approximately 30 min at room temperature and incubated with primary antibodies overnight at 4 °C. After washing, fluorescent secondary antibodies were applied for 1 h at room temperature in the dark. Nuclei were counterstained with DAPI, and slides were mounted with antifade mounting medium. Images were acquired using consistent exposure and acquisition settings across groups and quantified in ImageJ after background subtraction.

The antibodies used for tumor-tissue immunofluorescence included TNC (Servicebio, GB111243-100), TUBB3 (Servicebio, GB15139-100), CDH11/Cadherin-11 (Thermo Fisher Scientific, 32-1700), HAS3 (Thermo Fisher Scientific, PA5-76964), CEMIP (Bioss, bs-21528R), EGR1 (Proteintech, 12544-1-AP) and FOSL1 (Zhonghe, ZHS2608).

### Blood biochemistry, hematology, and ELISA

Blood was collected at the endpoint into anticoagulant and serum-separation tubes. Anticoagulated whole blood was used for routine hematology, including white blood cell count, lymphocyte percentage, red blood cell count, hemoglobin and platelet count. Serum was prepared from coagulated blood by centrifugation at 3,000 rpm for 10 min and used for biochemical assays, including total protein, alanine aminotransferase, aspartate aminotransferase, creatinine, blood urea nitrogen, uric acid and total bilirubin. Plasma for miRNA and ELISA assays was obtained from EDTA-anticoagulated blood after centrifugation at 3,000 rpm for 10 min at 4 °C and stored at −80 °C.

Plasma IL-6, TNF-α, IgM and IgG were measured using mouse ELISA kits according to the manufacturers’ instructions. Standards and plasma samples were equilibrated to room temperature before analysis. Each well received 50 μl standard or sample. After incubation at 37 °C, wells were washed three times, incubated with enzyme-conjugated secondary reagents, developed with substrate and stopped according to the kit protocol. Absorbance was read at 450 nm, and concentrations were calculated from standard curves.

### Statistics and reproducibility

Data are presented as mean ± s.d. or mean ± s.e.m. as specified in each figure legend. Biological replicates are defined in the corresponding Methods sections and figure legends. RT-qPCR assays used three biological replicates. Statistical analyses were performed using biological-replicate means. Technical replicates were averaged within each biological replicate before statistical analysis, and the statistical unit was the independent cell-culture experiment or individual mouse, as appropriate. RNA-seq used three biological replicates per group. Animal experiments used n = 3 mice per group and were interpreted as proof-of-concept in vivo experiments. Bar-graph comparisons among multiple treatment groups were performed using one-way ANOVA followed by Tukey’s multiple-comparisons test. For Fig. 6g and Fig. 7g, each plasma miRNA was compared between the gemcitabine and gemcitabine + M-PRIME groups using a two-sided unpaired Student’s t-test, with the individual mouse as the statistical unit. Unadjusted *p* values are shown in the corresponding panels, and *p* values from the three miRNA comparisons within each panel were additionally adjusted using the Holm–Šídák method. Effect sizes were reported as between-group mean differences with 95% confidence intervals. Longitudinal bioluminescence data were analyzed using two-way repeated-measures ANOVA followed by the Tukey’s multiple-comparisons test. Differential-expression analysis used adjusted *p* values to control for multiple testing. GraphPad Prism was used for plotting and statistical analysis of experimental data, and Python was used for transcriptomic data processing. All statistical tests were two-sided.

## Supporting information

This PDF file includes: Supplementary Figures 1-34 and Supplementary Table 1.

## Acknowledgements

This study was funded by the National Natural Science Foundation of China (grant numbers 22325801 to H.L. and 32271521 to X.S.) and Beijing Natural Science Foundation (grant numbers 26L080043 to H.L. and X.S.).

## Author Contributions

X.S. and H.L. conceived and supervised the study, secured funding and revised the manuscript. Z.Z. contributed to the conception and design of the study, designed and performed the experiments, analysed and interpreted the data, and drafted the manuscript. L.Z., X.W. and C.Z. contributed to the experiments and data analysis. Q.Y. and S.W. assisted with methodology development and interpretation of the results. All authors discussed the results and contributed to manuscript revision.

## Competing interests

The authors declare no competing interests.

## Data availability

The data supporting the plots in this paper are available from the corresponding author upon reasonable request.

## Code availability

The custom code used for data analysis in this study, including RNA-seq data processing, differential expression analysis, miRNA target prioritization, gene enrichment analysis and data visualization, is available at GitHub: https://github.com/Su-Laboratory/PRIME. Other analyses were performed using publicly available software as described in the Methods section.

## Supplementary information

Supplementary Table 1 and Supplementary Figs. 1-34.

## Notes

### Competing Interest Statement

The authors have declared no competing interest.

