## Supplementary material for "Sequence-specific and programmable catalytic degradation of extracellular microRNAs by membrane-anchored DNA nanomachines": This PDF file includes: Supplementary Figures 1-34 and Supplementary Table 1.

*Supplementary Figures 1-34.*

*Supplementary Table 1.*

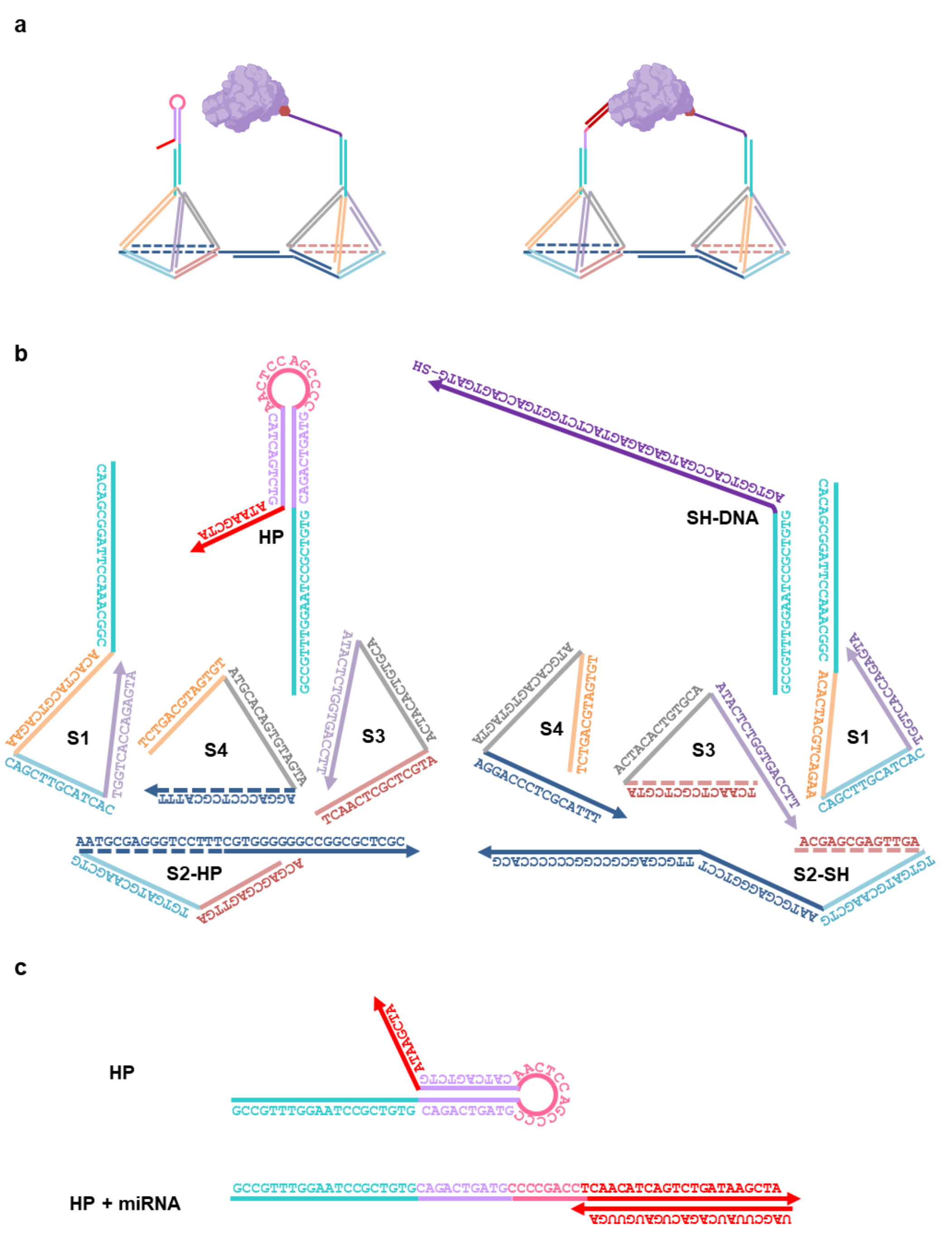


**Supplementary Fig. 1**: miRNA-responsive PRIME architecture and sequence design. a, Schematic of PRIME before and after target-miRNA binding. The double-tetrahedral scaffold positions a programmable stem-loop recognition unit near an RNase H-DNA loading unit. Target miRNA opens the hairpin to generate a local RNA/DNA hybrid for RNase H-mediated cleavage. b, DNA strands used to assemble PRIME, with colours matching the structural domains in panel a. c, miR-21-responsive hairpin design in the closed and target-bound states. Arrows indicate 5'-to-3' strand direction.

| | 1 | 2 | 3 | 4 | 5 | 6 | 7 | 8 |
|---|---|---|---|---|---|---|---|---|
| S1 | + | + | + | + | + | + | + | + |
| S2-HP | | + | + | + | | + | | + |
| S2-SH | | | | | + | | + | + |
| S3 | | | + | + | + | + | + | + |
| S4 | | | | + | + | + | + | + |
| HP | | | | | | + | | + |
| SH-DNA | | | | | | | + | + |

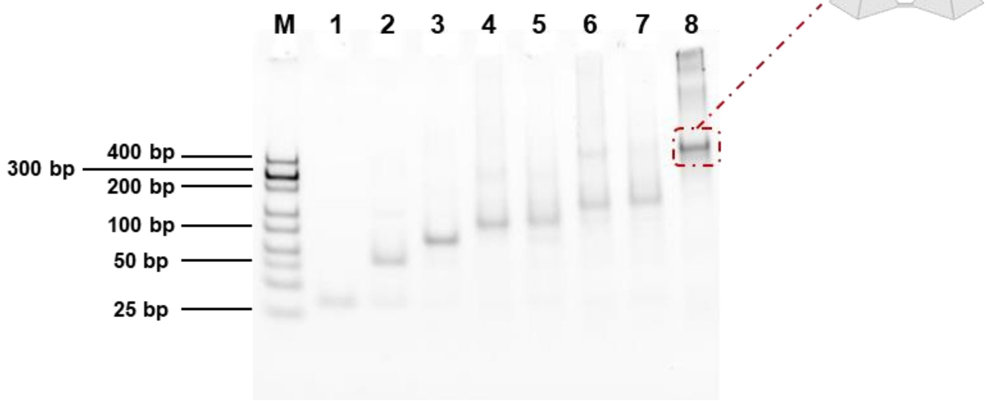


**Supplementary Fig. 2**: Stepwise native PAGE validation of the double-tetrahedral PRIME scaffold. DNA mixtures were assembled with the strand combinations indicated above the gel. Lanes 1–4 show stepwise assembly of one tetrahedral subunit, and lane 5 shows the second tetrahedral subunit. Lanes 6 and 7 show the two functionalized tetrahedral units bearing the stem-loop recognition domain and RNase H-recruiting strand, respectively. Lane 8 shows the double-tetrahedral PRIME scaffold formed by bridge-domain hybridization. M, DNA marker. Electrophoresis was performed on a 4-20% gradient native PAGE gel at 80 V for 50 min.

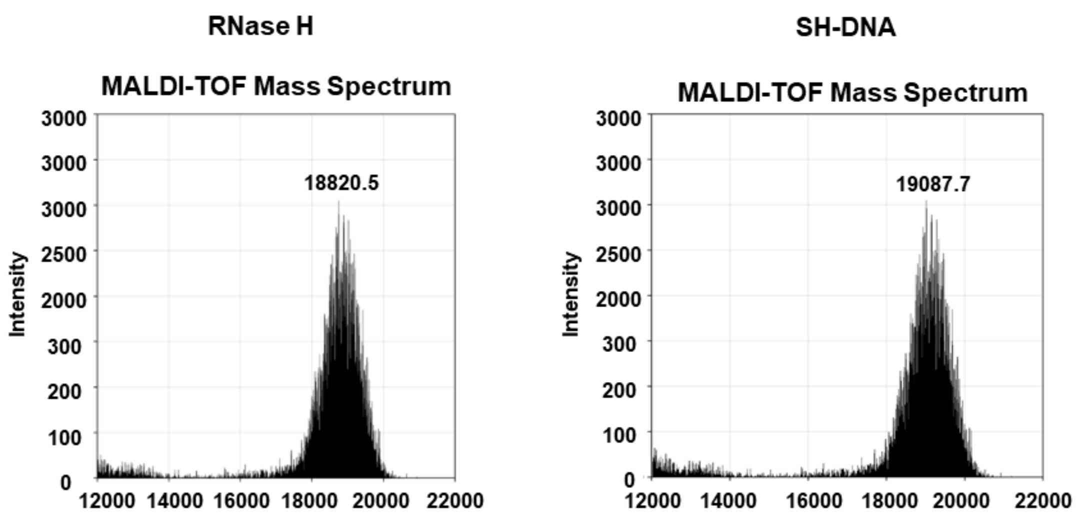


**Supplementary Fig. 3**: MALDI-TOF evidence for protein-DNA conjugation. MALDI-TOF mass spectra of RNase H and thiol-modified SH-DNA used to generate the catalytic loading unit. RNase H showed an observed mass peak at 18,820.5 m/z, and SH-DNA showed an observed mass peak at 19,087.7 m/z. Together with the conjugate peak in Fig. 2b, these spectra support coupling of RNase H to SH-DNA.

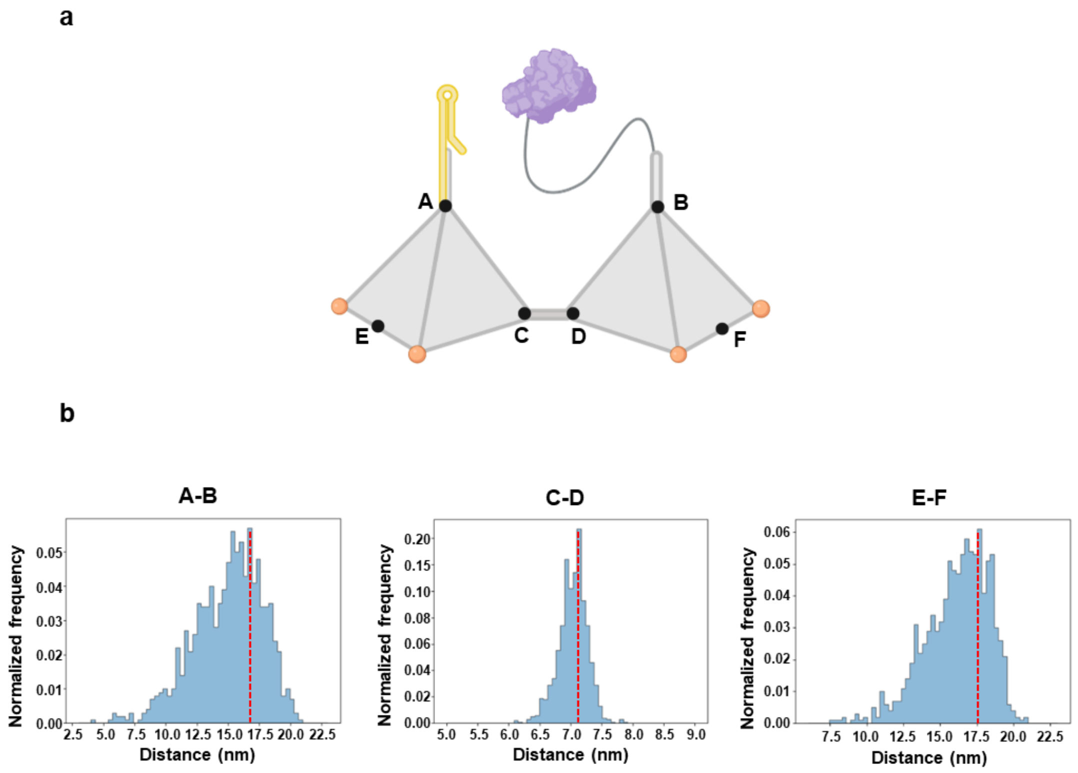


**Supplementary Fig. 4**: oxDNA distance analysis of PRIME scaffold geometry. a, Schematic of PRIME showing the positions used for oxDNA distance measurements.

Black spheres mark nucleotide positions selected for intramolecular distance measurements. b, Normalized distance distributions for A-B, C-D and E-F position pairs across the oxDNA trajectory. Red dashed lines indicate mean distances. Distances are reported in nanometres.

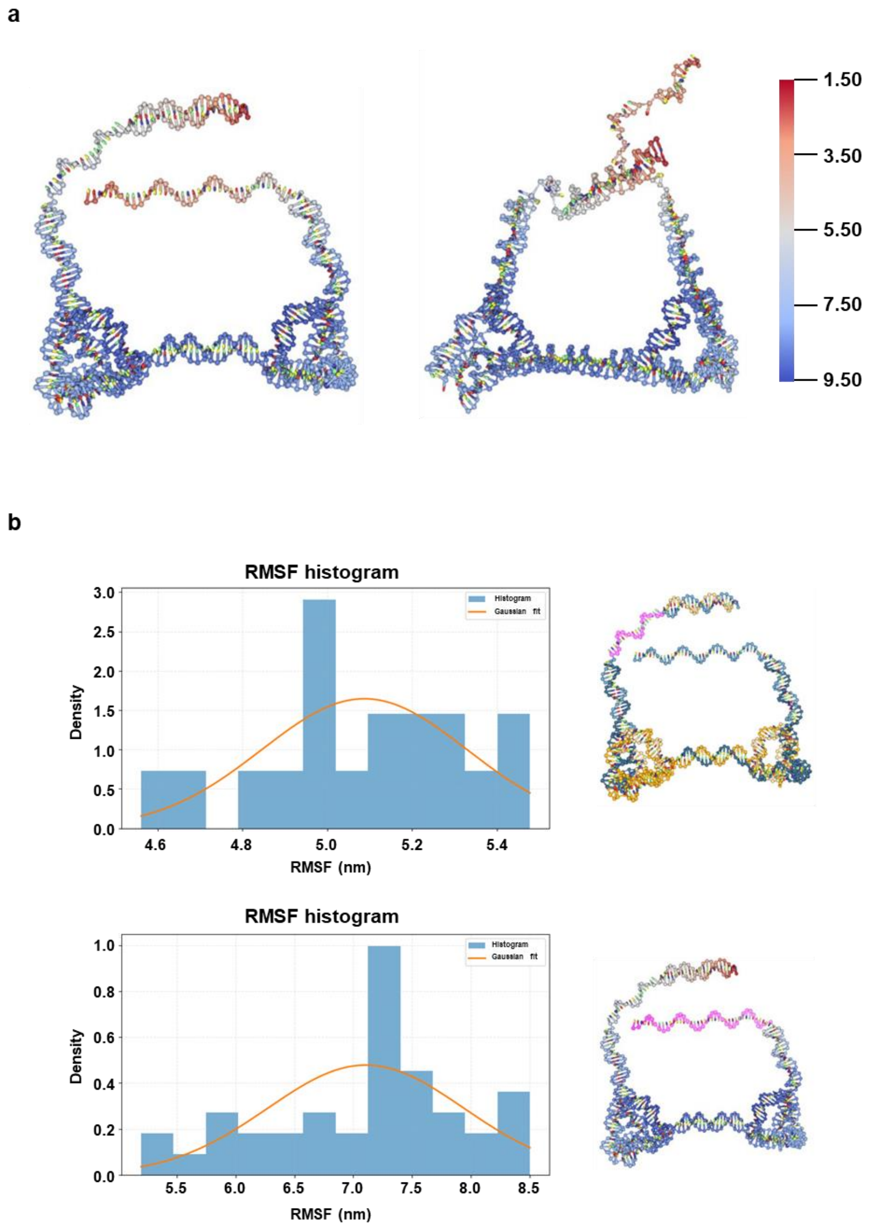


**Supplementary Fig. 5**: Simulated flexibility of the double-tetrahedral PRIME scaffold. a, Representative oxDNA conformations viewed from different orientations and colour-coded by root mean square fluctuation (RMSF). b, RMSF distributions for flexible segments near the substrate-recognition and RNase H-loading regions. Blue bars show the RMSF distributions, orange curves indicate Gaussian fits, and the corresponding analysed segments are highlighted in magenta on the structural models. Local flexibility in these regions may facilitate substrate access within the scaffold. RMSF values are reported in nanometres.

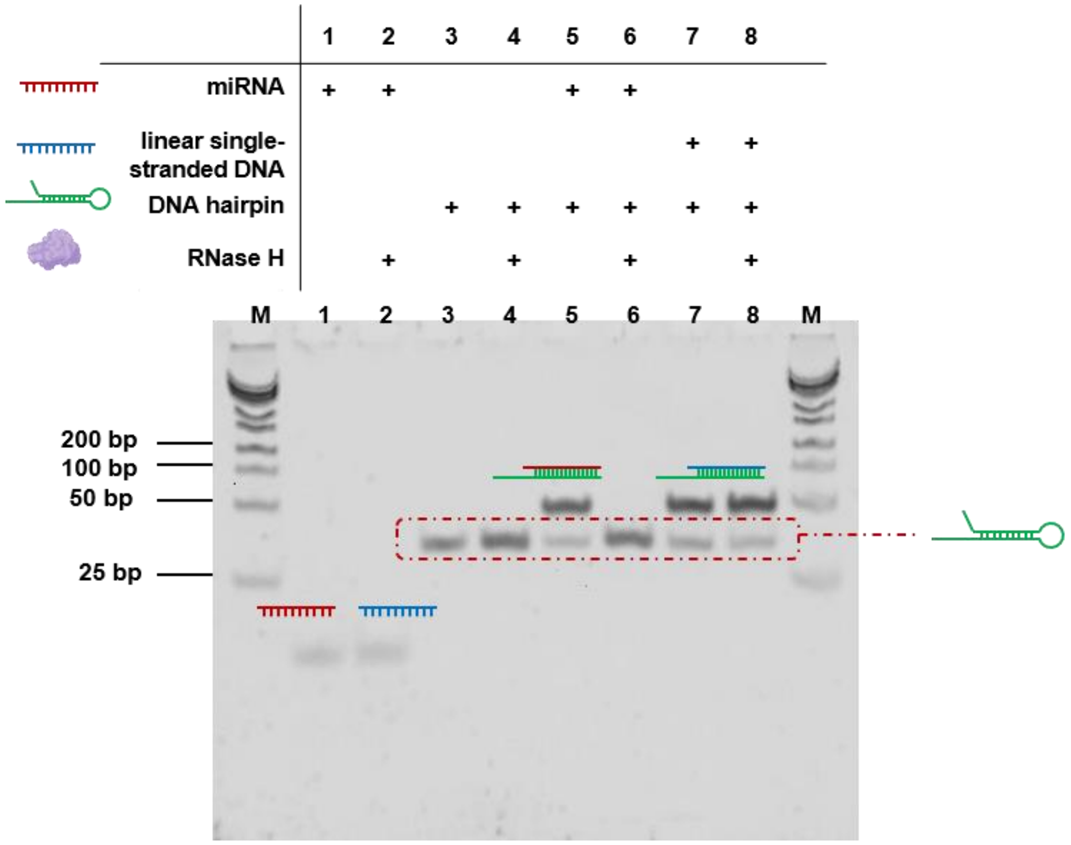


**Supplementary Fig. 6**: RNase H selectively cleaves RNA/DNA hybrid substrates. Native PAGE analysis of miR-21 (lanes 1 and 2), the DNA hairpin (lanes 3 and 4), the miR-21/DNA hairpin hybrid (lanes 5 and 6) and the linear DNA/DNA hairpin hybrid (lanes 7 and 8). RNase H was added to the even-numbered lanes. Cleavage was detected only for the RNA/DNA hybrid substrate, supporting RNase H-dependent degradation of target miRNA after hairpin opening. M, DNA marker. Electrophoresis was performed on a 10% native PAGE gel at 120 V for 40 min.

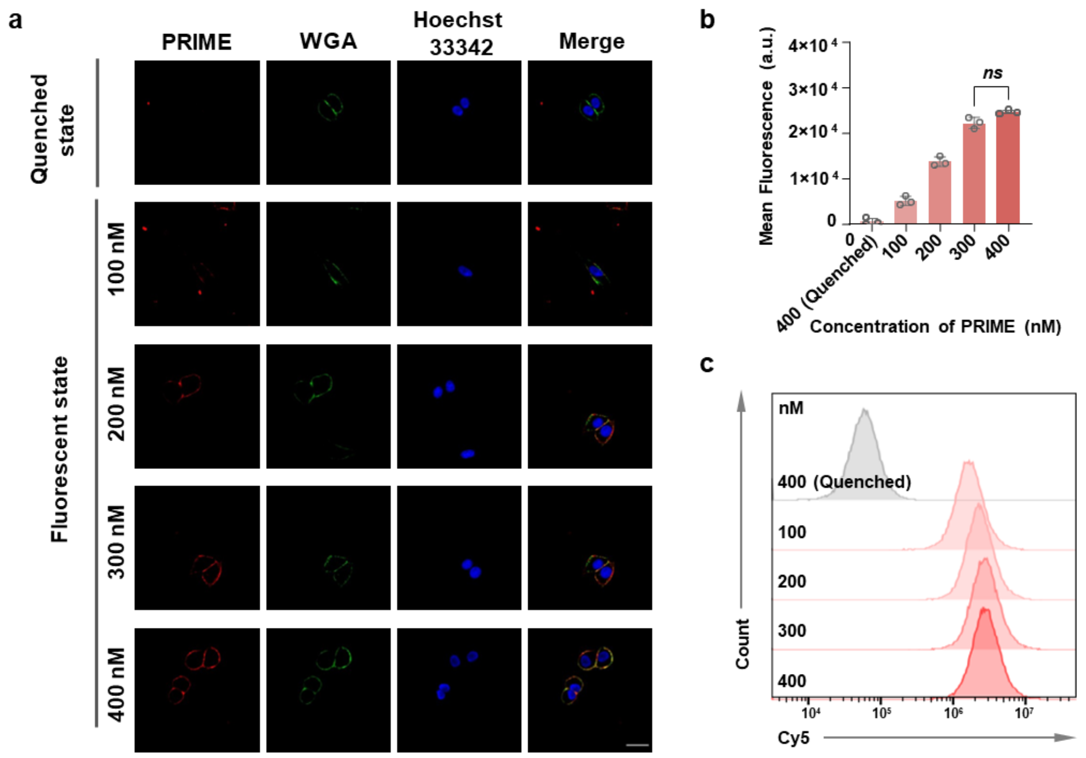


**Supplementary Fig. 7**: Concentration optimization of cholesterol-anchored PRIME for membrane loading. a, Representative fluorescence images of cells incubated with Cy5-labelled cholesterol-modified unquenched PRIME at 100, 200, 300 or 400 nM. Images show PRIME, wheat germ agglutinin (WGA)-labelled plasma membrane, Hoechst-stained nuclei and merged channels. A quenched PRIME sample served as a fluorescence-background control. Scale bar, 10 µm. b, Quantification of membrane-associated fluorescence intensity. Data are mean ± s.d. from three independent experiments. c, Flow-cytometry histograms of Cy5 fluorescence after PRIME loading. For each sample, 10,000 events were collected.

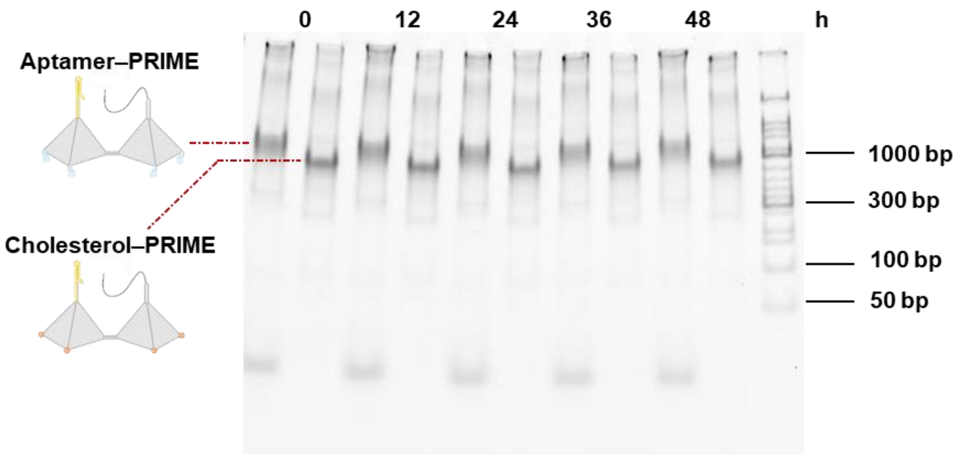


**Supplementary Fig. 8**: Serum stability of aptamer- and cholesterol-modified PRIME scaffolds. Native PAGE analysis of PRIME nucleic-acid scaffolds after incubation in cell-culture medium containing 10% fetal bovine serum for the indicated durations. At each time point, the left and right lanes contain aptamer-modified and cholesterol-modified PRIME, respectively. M, DNA marker. Electrophoresis was performed on a 4-20% gradient native PAGE gel at 80 V for 50 min.

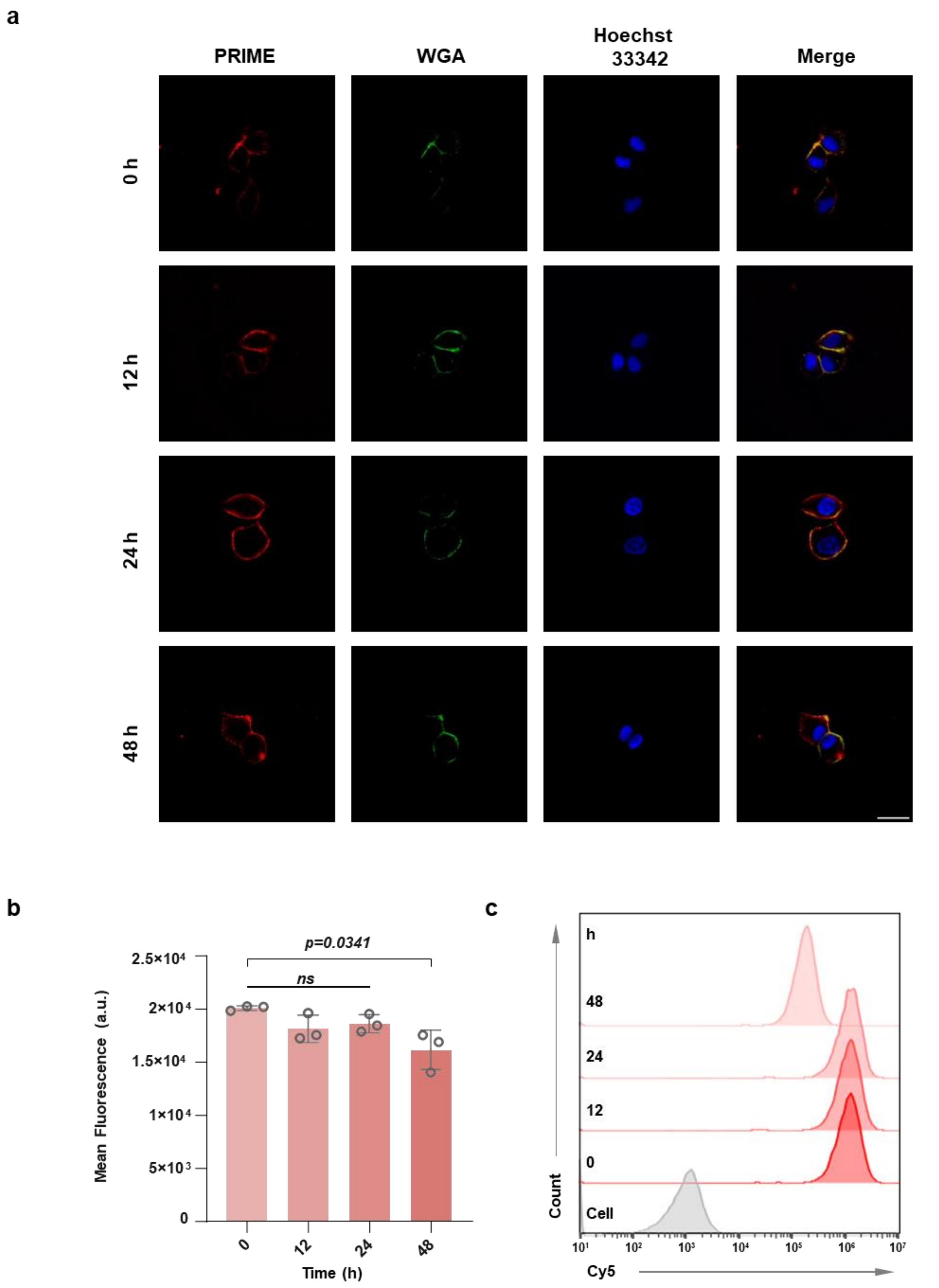


**Supplementary Fig. 9**: Persistence of cell-associated fluorescence after cholesterol-anchored PRIME loading. a, Representative fluorescence images of cells labelled with Cy5-labelled cholesterol-modified unquenched PRIME and imaged at 0, 12, 24 and 48 h after membrane insertion. Images show PRIME, WGA-labelled plasma membrane, Hoechst-stained nuclei and merged channels. Scale bar, 10 μm. b, Quantification of membrane-associated PRIME fluorescence over time. Data are mean ± s.d. from three independent experiments. c, Flow-cytometry histograms of Cy5 fluorescence from labelled cells; unlabeled cells served as background controls. For each sample, 10,000 events were collected.

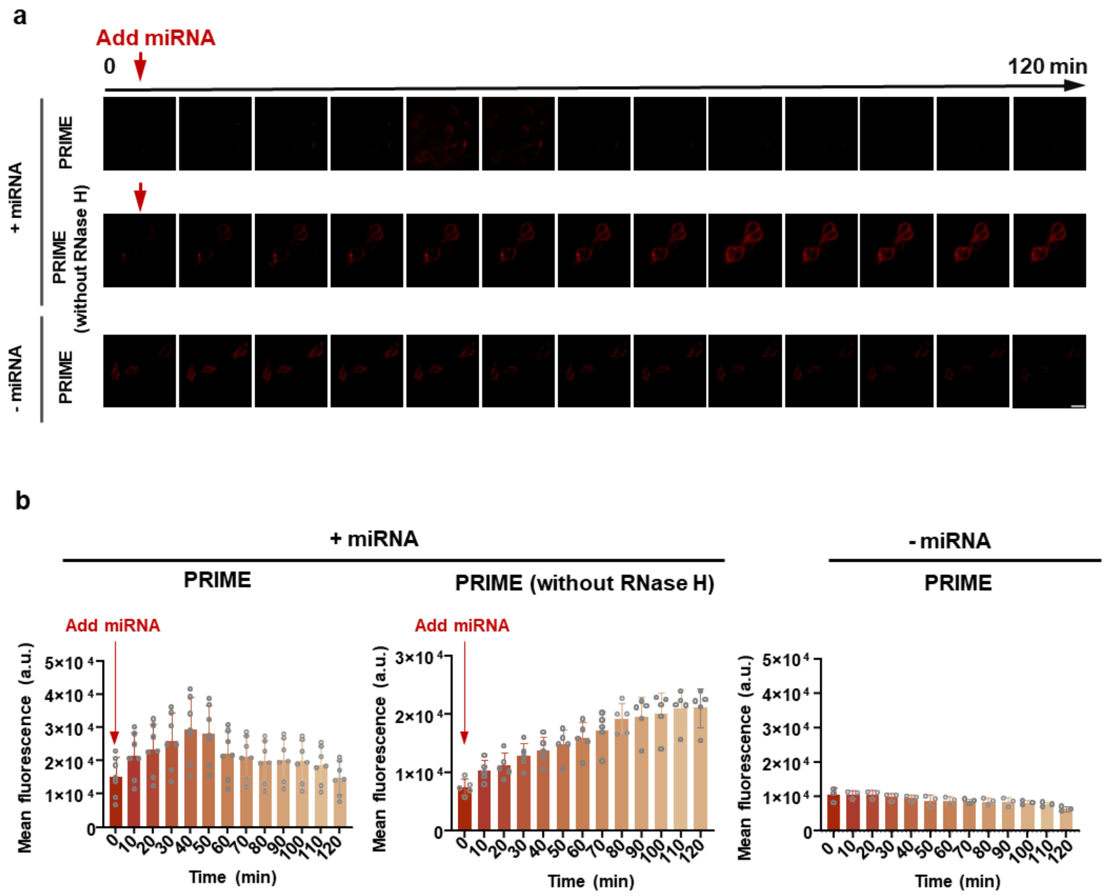


**Supplementary Fig. 10**: Imaging controls for membrane-anchored PRIME reaction cycles. a, Representative time-lapse TIRF images of cells labelled with fluorophore-quencher-labelled cholesterol-modified PRIME. Red arrows mark miRNA addition. Complete PRIME with miR-21 shows activation followed by decay, RNase H-deficient PRIME shows a sustained fluorescence increase, and complete PRIME without added miRNA remains near baseline. Scale bar, 10 μm. b, Quantification of membrane-associated fluorescence over time. Data are mean ± s.d. from at least three biological replicates.

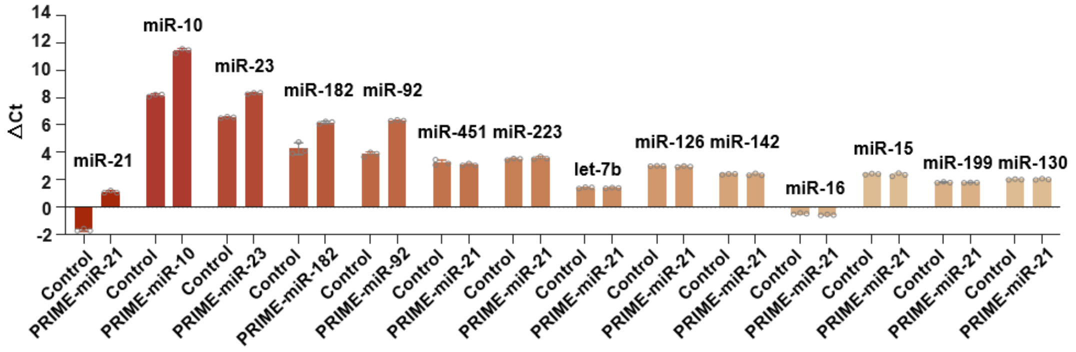


**Supplementary Fig. 11**: Raw ΔCt values for RT-qPCR analysis corresponding to Fig. 4d. Extracellular miRNAs in culture supernatants were measured after treatment with the indicated sequence-programmed PRIME formulations. Group labels indicate the PRIME formulation and the miRNA quantified. The y axis shows ΔCt values calculated as Ct_target miRNA - Ct_spike-in control before conversion to relative extracellular miRNA abundance in Fig. 4d. Data are mean ± s.d. from three biological replicates.

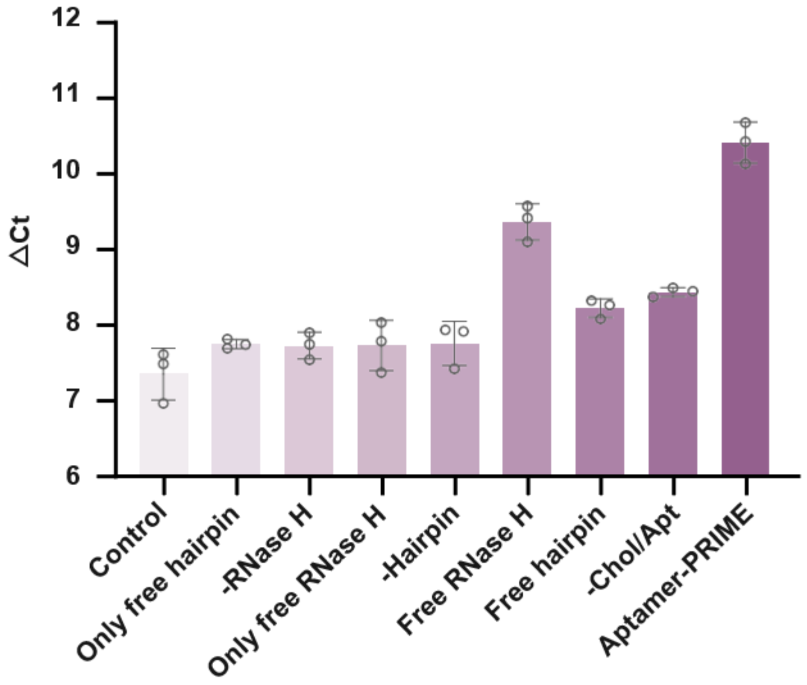


**Supplementary Fig. 12**: Raw ΔCt values for RT-qPCR analysis corresponding to Fig. 4e. Extracellular miR-21 in culture supernatants was measured after treatment with the indicated PRIME-miR-21 variants. The y axis shows ΔCt values calculated as Ct_miR-21 - Ct_spike-in control before conversion to relative extracellular miR-21 abundance in Fig. 4e. The Chol-PRIME group in Fig. 4e uses the same RT–qPCR dataset as the first group in Fig. 4d (Supplementary Fig. 11) and is therefore not replotted here. Data are mean ± s.d. from three biological replicates, each measured in technical triplicate. Each plotted point represents the mean of the three technical replicates from one biological replicate.

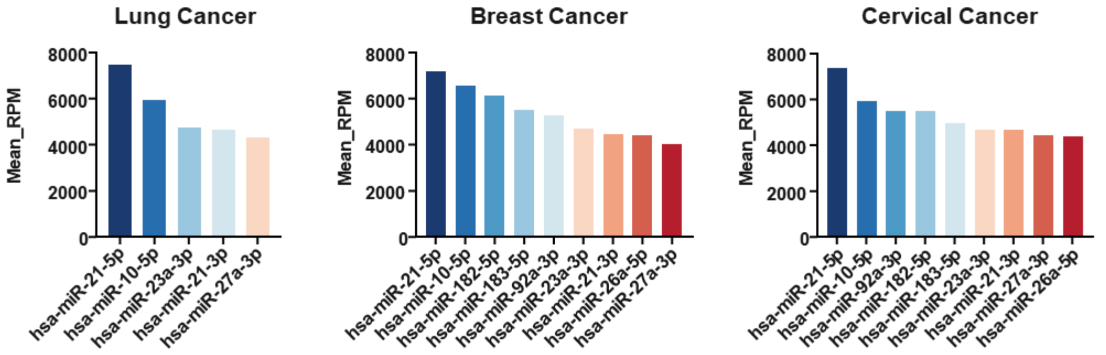


**Supplementary Fig. 13**: Dataset-guided prioritization of extracellular pro-metastatic miRNA targets. For GEO evidence, 36 NSCLC plasma profiles from 18 patients in GSE244643 were used, whereas the HNSCC samples in this dataset were not included. All 12 breast-cancer plasma samples in GSE240872 were included. For GSE302097, which contains plasma, urine and liquid-based cytology (LBC) samples from subjects with high-grade cervical intraepithelial lesions, only the 24 plasma samples were used. For TCGA tumour–normal classification, the processed matrices comprised 450 tumour and 45 normal LUAD samples, 749 tumour and 76 normal BRCA samples, and 306 tumour and 3 normal CESC samples. Candidate miRNAs were identified by integrating TCGA mature-miRNA abundance, GEO-derived evidence of extracellular occurrence and migration- or metastasis-associated annotations, and were then ranked within each cancer context in descending order of mean TCGA mature-miRNA RPM. Bar heights represent mean TCGA mature-miRNA abundance expressed as reads per million mapped mature-miRNA reads (RPM); GEO-derived values were used only as extracellular-miRNA evidence and were not used as the plotted quantitative measure. Mature miRNAs are labelled using their full miRBase nomenclature, including the species prefix and mature-arm designation (for example, hsa-miR-21-5p). The selected target sets were subsequently used to program PRIME formulations for lung, breast and cervical cancer cell models; this prioritization reflects extracellular occurrence, literature-supported migration or metastasis association and TCGA abundance, rather than a direct measurement of pro-metastatic activity.

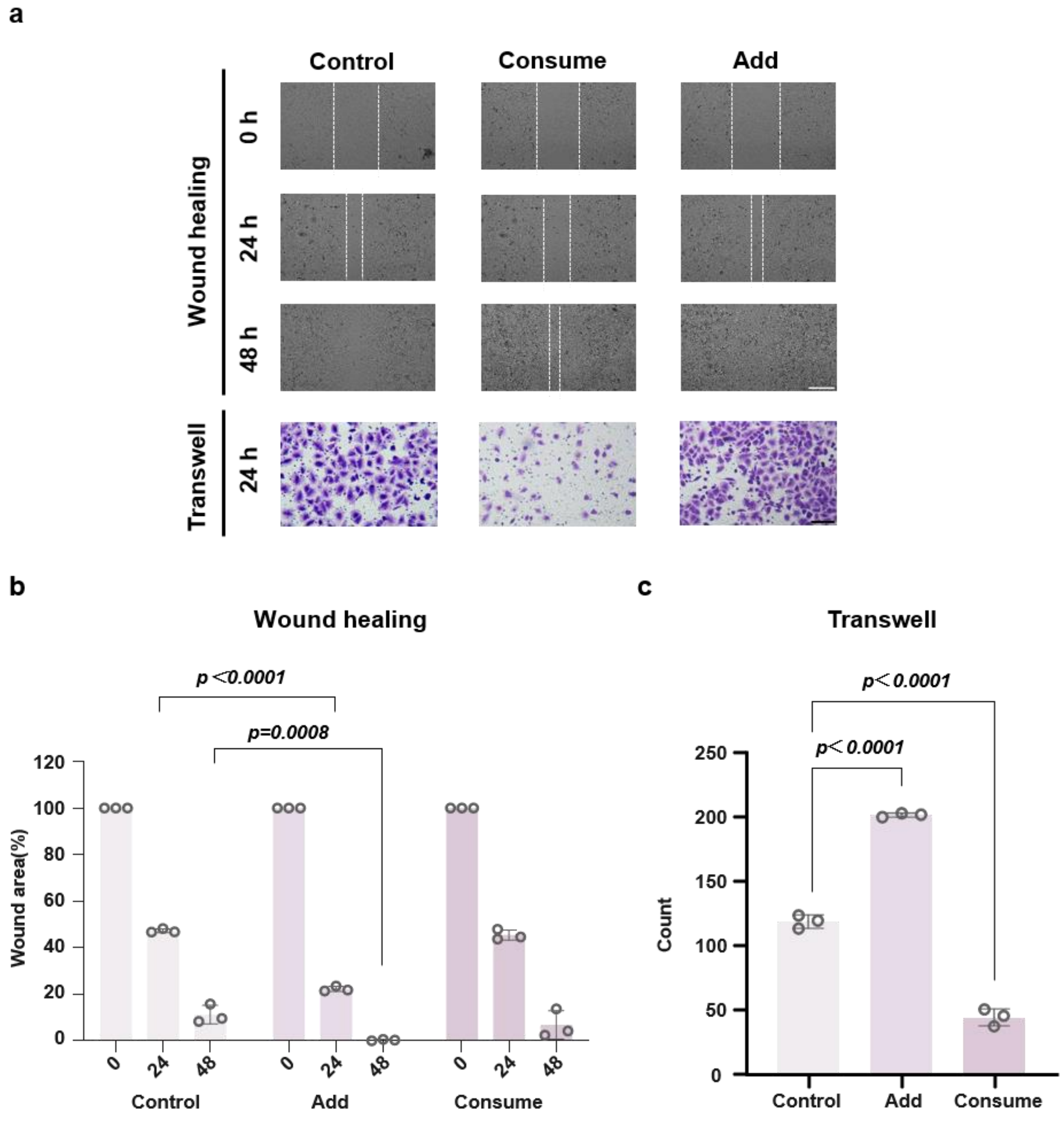


**Supplementary Fig. 14**: PRIME-miR-21 attenuates exogenous miR-21-induced migration. a, Representative wound-healing and Transwell images of A549 cells after extracellular miR-21 supplementation. Add denotes supplementation with exogenous miR-21, whereas Consume denotes cells pre-modified with cholesterol-PRIME-miR-21 before the same exogenous miR-21 supplementation. For wound-healing assays, synthetic miR-21 was added at a final concentration of 100 nM in 2 ml of medium per well (200 pmol per well). For Transwell assays, miR-21 was added at a final concentration of 100 nM to both the upper chamber (200 µl; 20 pmol) and lower chamber (500 µl; 50 pmol). Wound-healing images were acquired at 0, 24 and 48 h, and Transwell images after 24 h. b,c, Quantification of remaining wound area and migrated-cell counts. Data are mean ± s.d. from independent experiments. Representative images are shown with scale bars of 100 µm for wound-healing assays and 20 µm for Transwell assays.

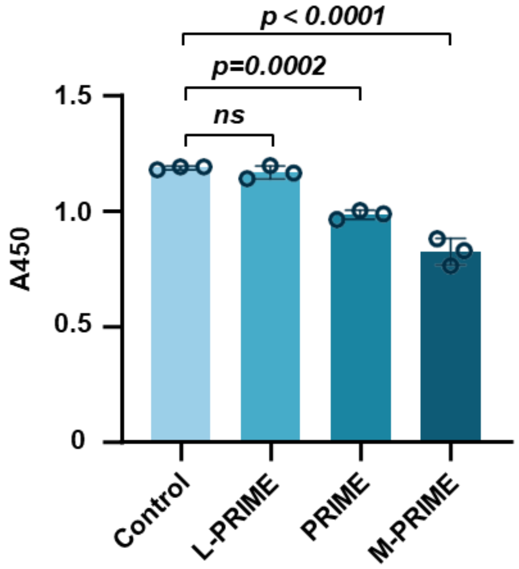


| | Control | L-PRIME | PRIME | M-PRIME |
|---|---|---|---|---|
| Inhibition rate | 0 | 1.9% | 19.4% | 34.4% |

**Supplementary Fig. 15**: Metabolic-activity control for A549 cells treated with PRIME formulations. CCK-8 absorbance and signal inhibition rates were measured after treatment with 100 nM PRIME for 48 h. L-PRIME denotes the control mixture targeting the low-ranked miRNAs miR-4438, miR-4665 and miR-3686, whereas M-PRIME targets the high-ranked miRNAs miR-21, miR-10 and miR-23. The inhibition rate was calculated as $[(OD_{control} - OD_{treated})/OD_{control}] \times 100\%$. Data are mean ± s.d. from three independent experiments.

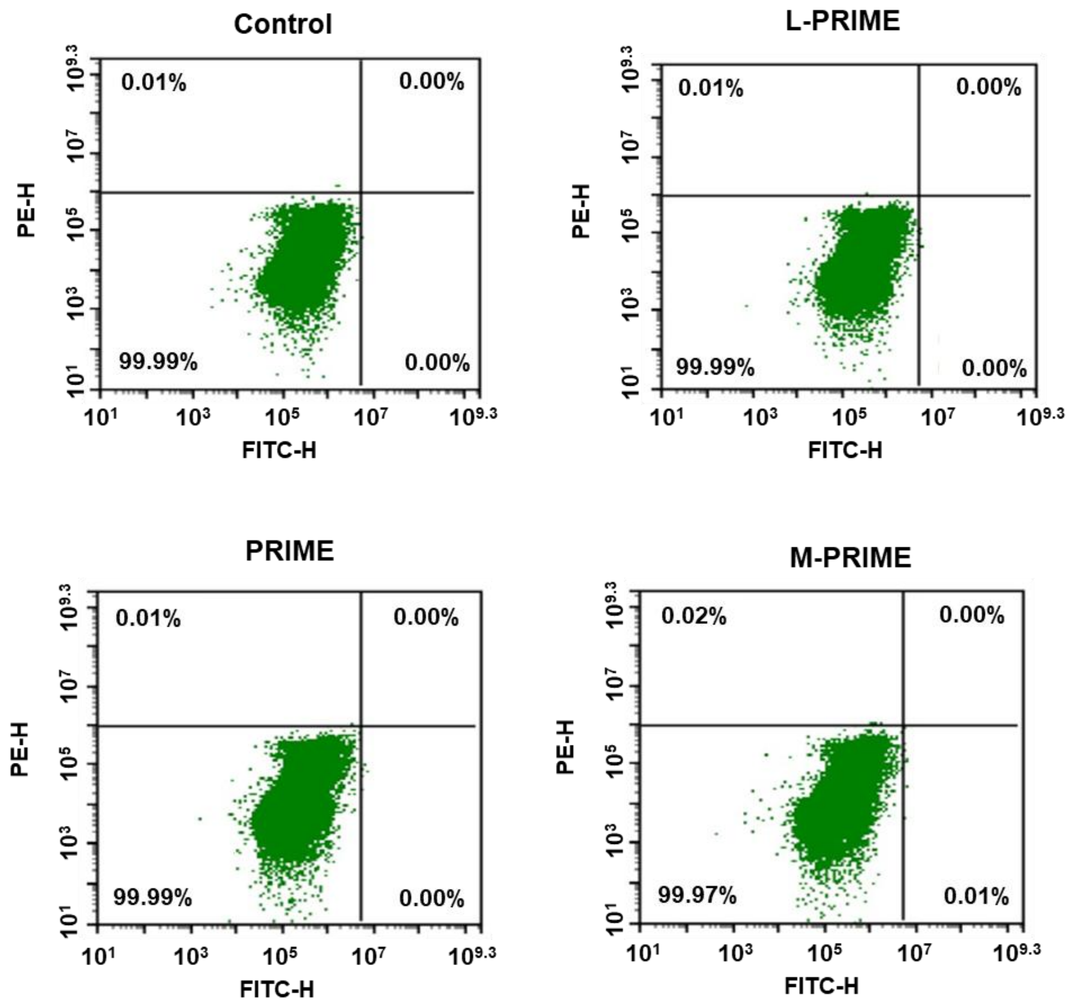


**Supplementary Fig. 16**: Apoptosis analysis after PRIME treatment. Compensation-adjusted Annexin V/PI flow-cytometry plots of A549 cells after treatment with the indicated PRIME formulations at 100 nM for 48 h. For each sample, 10,000 events were collected. Quadrants indicate viable cells, early apoptotic cells, late apoptotic or secondary necrotic cells, and PI-positive dead cells.

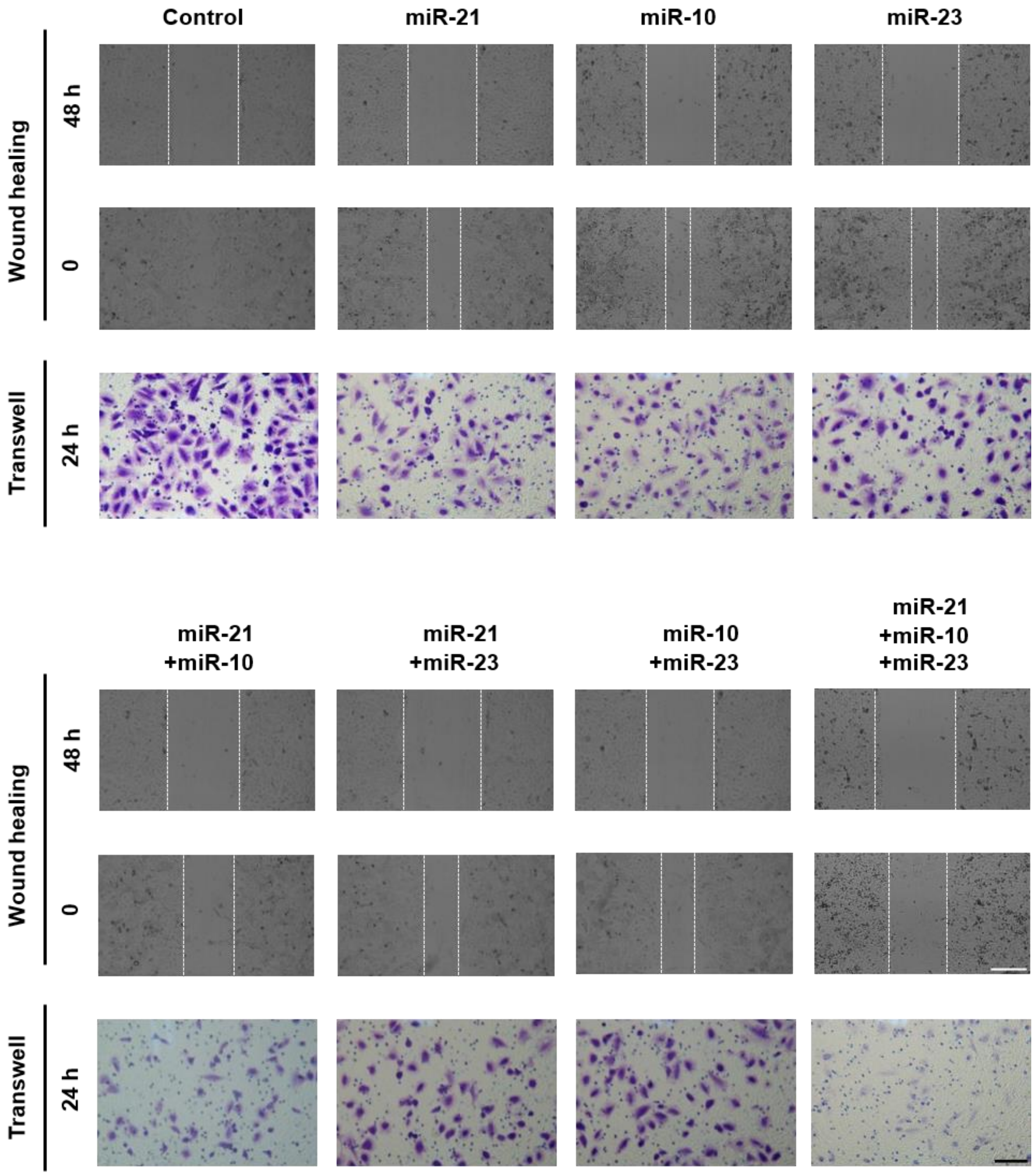


**Supplementary Fig. 17**: Single-, dual- and triple-target PRIME treatment of A549 migration. Representative wound-healing and Transwell images of A549 cells treated with cholesterol-modified PRIME constructs targeting the indicated lung-cancer-associated extracellular miRNAs. Targets were selected from the high-ranked lung cancer set in Supplementary Fig. 13. Wound-healing images were acquired at 0 and 48 h, and Transwell images after 24 h. Cells were serum-starved before both assays. Representative images are shown with scale bars of 100 μm for wound-healing assays and 20 μm for Transwell assays.

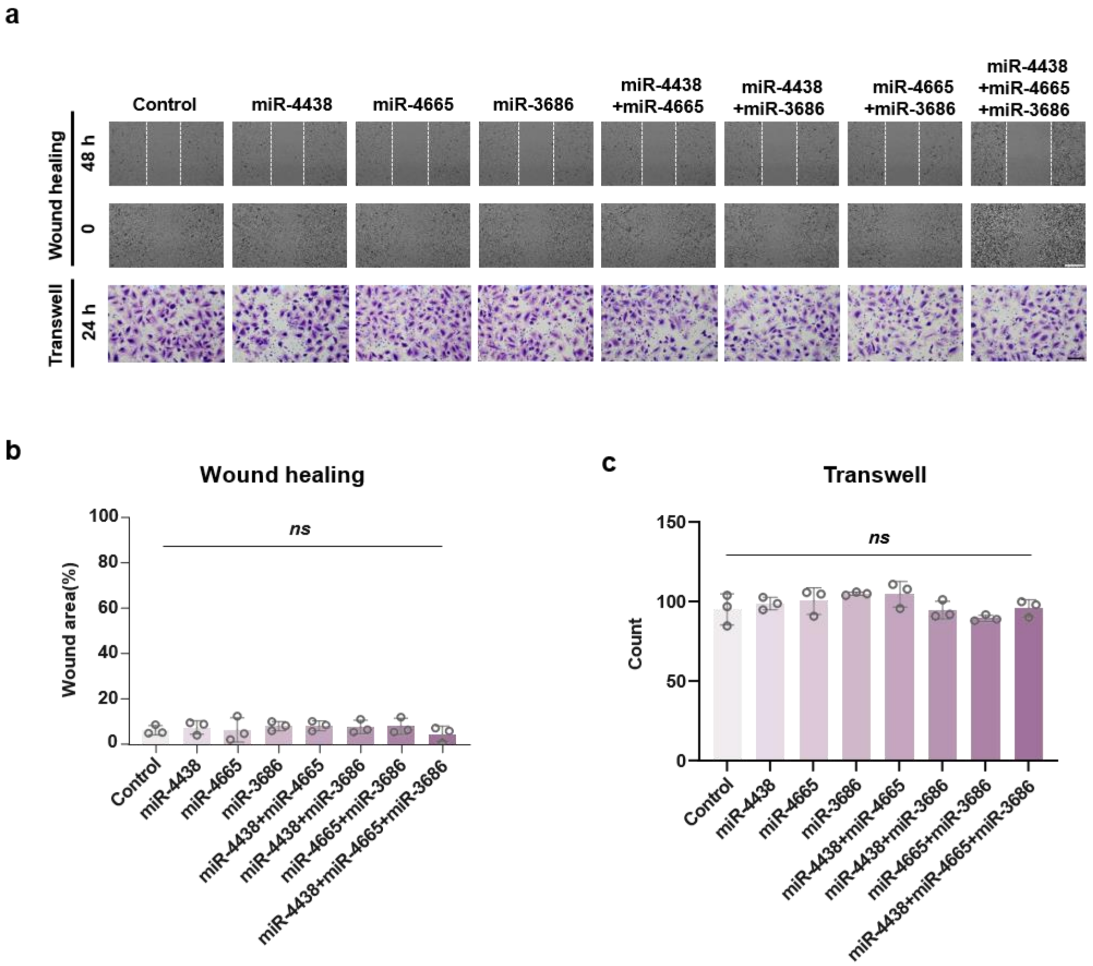


**Supplementary Fig. 18**: Targeting low-ranked extracellular miRNAs does not reproduce the anti-migratory effect of M-PRIME. a, Representative wound-healing and Transwell images of A549 cells treated with cholesterol-modified PRIME constructs targeting the low-ranked extracellular miRNAs miR-4438, miR-4665 and miR-3686, individually or in the indicated combinations. b,c, Quantification of remaining wound area and migrated-cell counts. Data are mean ± s.d. from independent experiments. Representative images are shown with scale bars of 100 μm for wound-healing assays and 20 μm for Transwell assays.

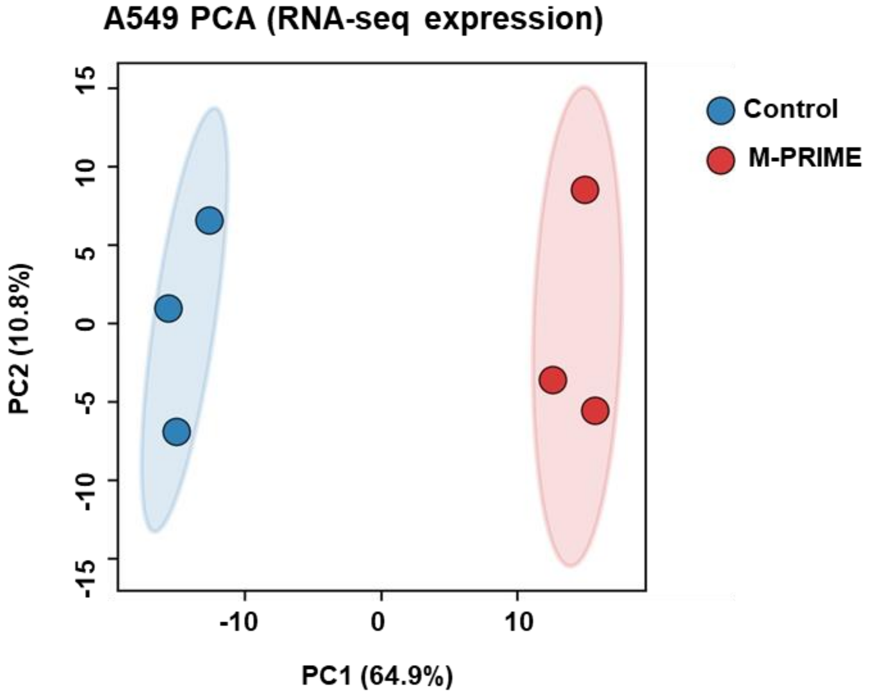


**Supplementary Fig. 19**: M-PRIME remodels the A549 transcriptomic state. Principal component analysis of RNA-seq profiles from A549 cells treated for 48 h with cholesterol-modified M-PRIME targeting miR-21, miR-10 and miR-23. Blue points indicate controls, red points indicate M-PRIME-treated samples and shaded ellipses indicate replicate distributions.

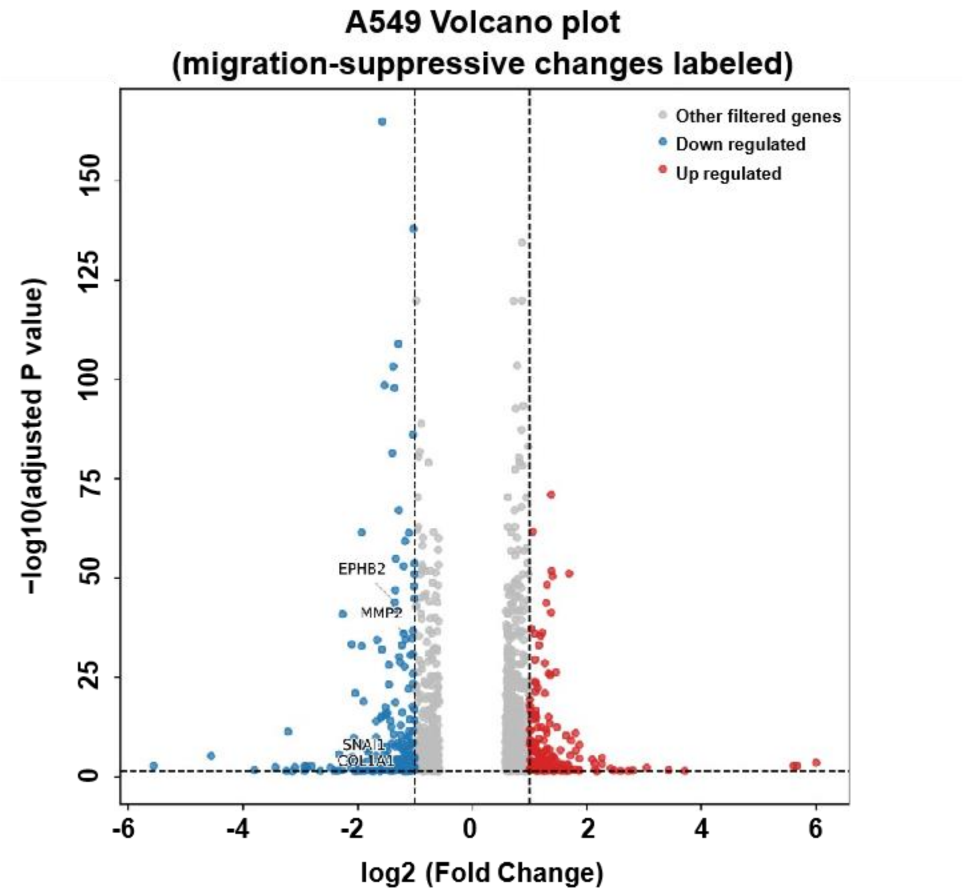


**Supplementary Fig. 20**: Differential-expression analysis after M-PRIME treatment. Volcano plot of RNA-seq changes in A549 cells after 48 h treatment with M-PRIME targeting miR-21, miR-10 and miR-23. Each point represents one gene. The x axis shows log2(fold change), and the y axis shows −log10(adjusted *p* value). Red and blue points indicate genes with log2(fold change) $\geq 1$ and adjusted $p < 0.05$ or log2(fold change) $\leq -1$ and adjusted $p < 0.05$, respectively, whereas grey points indicate the remaining genes. Vertical dashed lines mark log2(fold change) = ±1, and the horizontal dashed line marks −log10(0.05) = 1.301, corresponding to adjusted $p = 0.05$. Representative migration-associated downregulated genes are labelled.

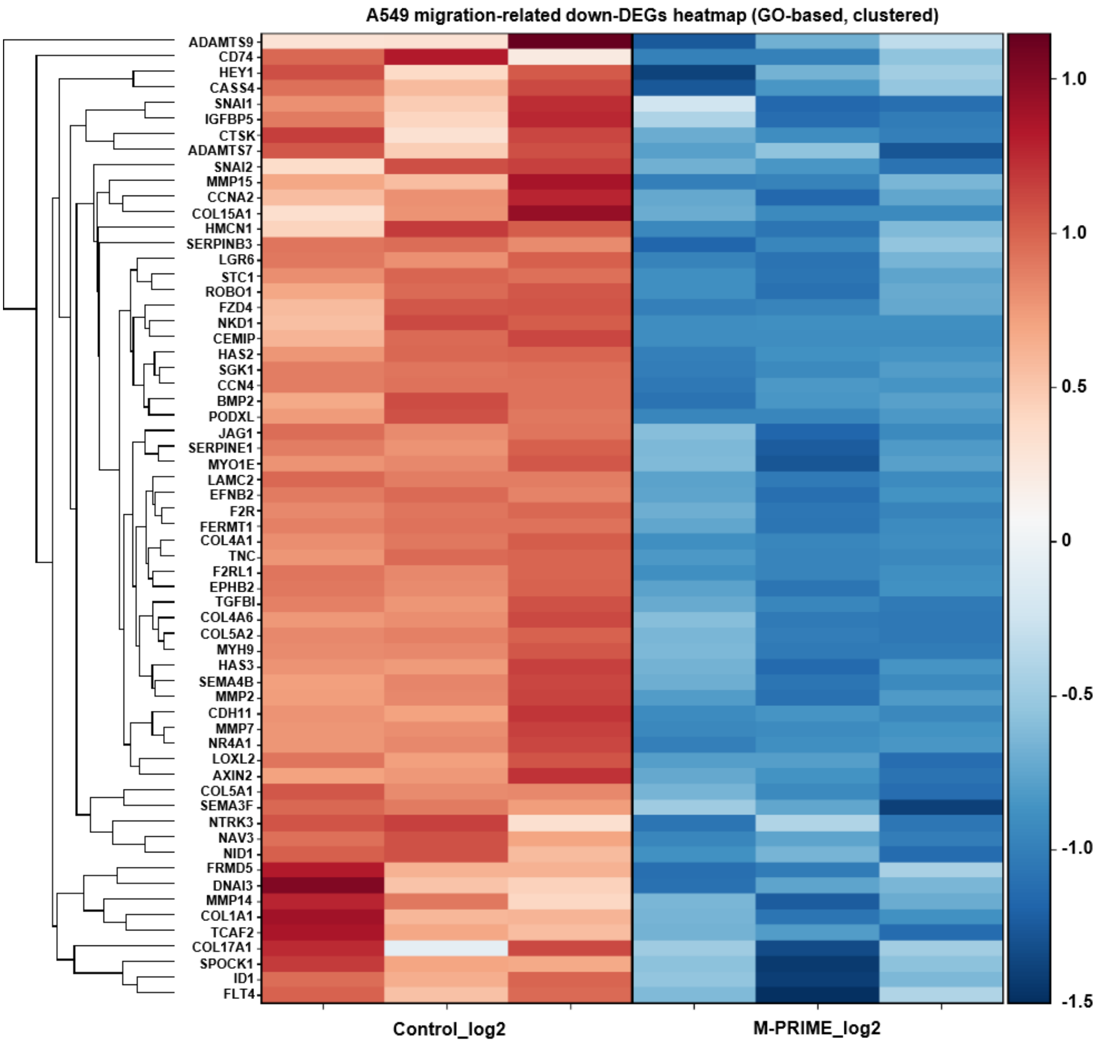


**Supplementary Fig. 21**: Downregulation of migration-associated gene programs after M-PRIME treatment. Clustered heat map of migration-associated downregulated genes in A549 cells treated for 48 h with cholesterol-modified M-PRIME targeting miR-21, miR-10 and miR-23. RNA-seq was performed using three biological replicates per group. Rows represent genes and columns represent the three control (ct1–ct3) and three M-PRIME-treated (tr1–tr3) biological replicates. Genes were selected by intersecting downregulated differentially expressed genes (log2(fold change) $\leq -0.58$ and adjusted $p < 0.05$) with genes contributing to significantly enriched migration-, motility-, adhesion-, cytoskeleton-, extracellular-matrix-, epithelial–mesenchymal transition- or invasion-related Gene Ontology terms (adjusted $p < 0.05$). The resulting genes were ranked by adjusted $p$ value and log2(fold change) before visualization and clustered using Euclidean distance and average linkage. Expression values are row-wise Z-scores calculated across the six biological samples from precomputed log2(FPKM + 1)-transformed expression values.

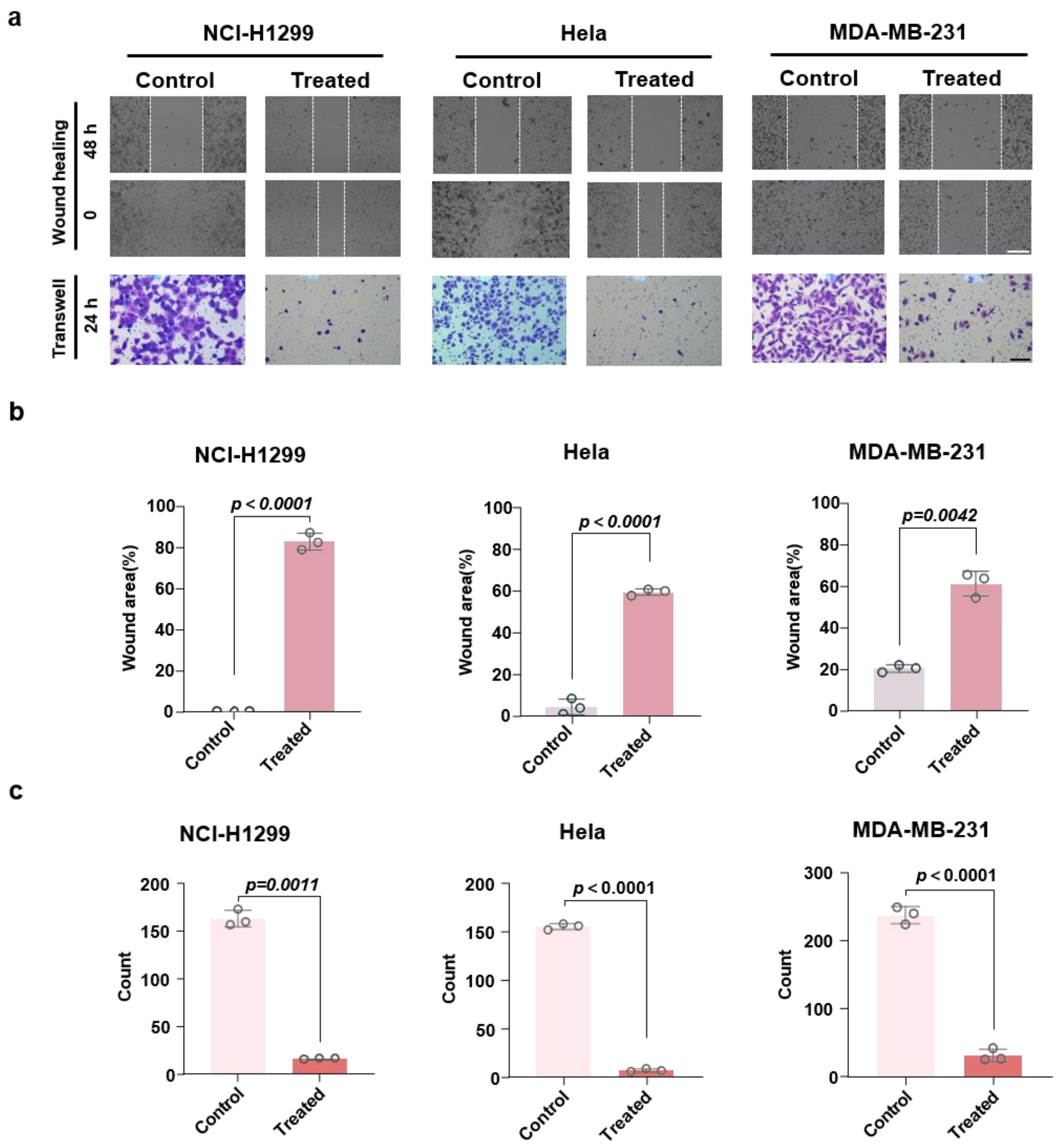


**Supplementary Fig. 22**: M-PRIME suppresses migration across additional tumour-cell models. a, Representative wound-healing and Transwell images of NCI-H1299 lung cancer cells, HeLa cervical cancer cells and MDA-MB-231 breast cancer cells treated with matched cholesterol-modified M-PRIME formulations. Target sets were miR-21/miR-10/miR-23 for lung cancer, miR-21/miR-10/miR-182 for breast cancer and miR-21/miR-10/miR-92 for cervical cancer. b,c, Quantification of remaining wound area and migrated-cell counts. Data are mean ± s.d. from independent experiments. Representative images are shown with scale bars of 100 μm for wound-healing assays and 20 μm for Transwell assays.

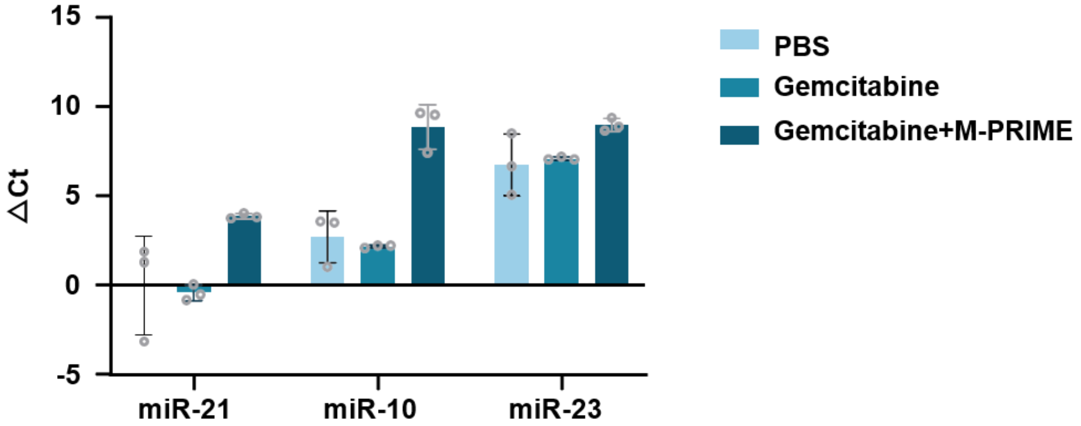


**Supplementary Fig. 23**: Raw ΔCt values for plasma miRNA RT-qPCR analysis corresponding to Fig. 6g. Plasma miR-21, miR-10 and miR-23 were measured in the orthotopic H1299-Luc lung-tumour model after treatment with PBS, gemcitabine or gemcitabine/M-PRIME. The y axis shows ΔCt values calculated as Ct_target miRNA - Ct_spike-in control before conversion to relative plasma miRNA abundance in Fig. 6g. Data are mean ± s.d. from plasma samples of three mice per group, each measured in technical triplicate. Each plotted point represents the mean of the three technical replicates from one mouse.

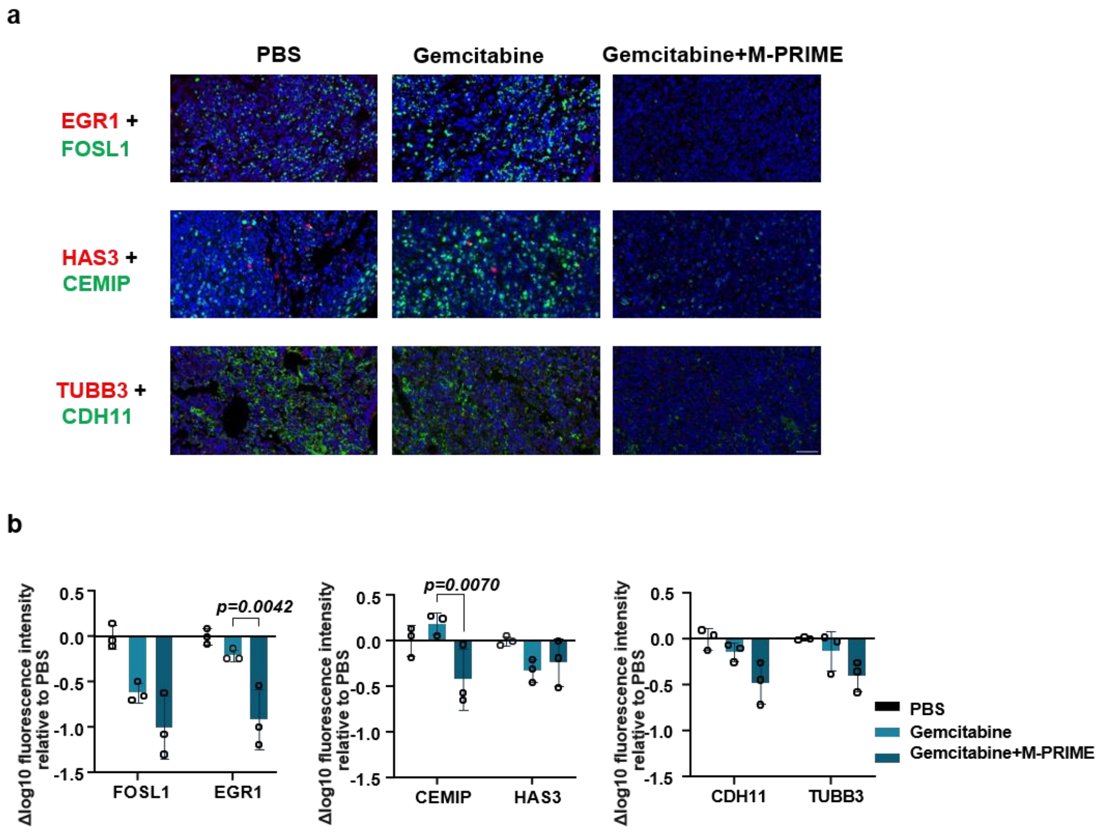


**Supplementary Fig. 24**: Tumour-tissue immunofluorescence after orthotopic lung-tumour treatment. a, Representative immunofluorescence images of tumour sections from mice treated with PBS, gemcitabine or gemcitabine/M-PRIME. Sections were stained for EGR1 and FOSL1, HAS3 and CEMIP, or TUBB3 and CDH11; nuclei were stained with DAPI. b, Quantification of background-corrected fluorescence intensities within tumour regions, log10-transformed and normalized as Δlog10 relative to PBS. Data are mean ± s.e.m. (n = 3 mice per group for quantitative analyses).

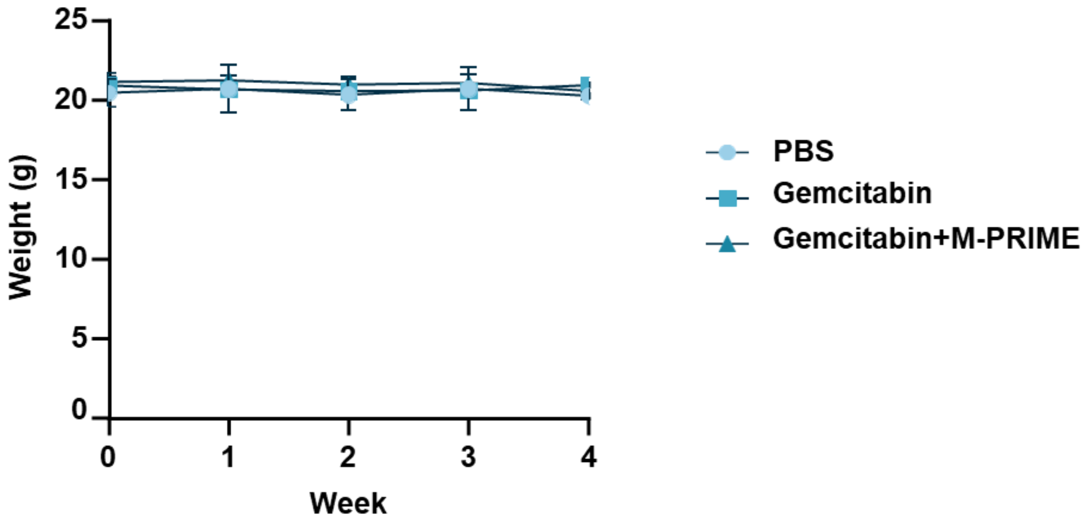


**Supplementary Fig. 25**: Body-weight monitoring in the orthotopic H1299-Luc lung-tumour model. Weekly body-weight changes of mice treated with PBS, gemcitabine or gemcitabine/M-PRIME. Body weight was recorded once per week throughout treatment. Data are mean ± s.d. (n = 3 mice per group for quantitative analyses).

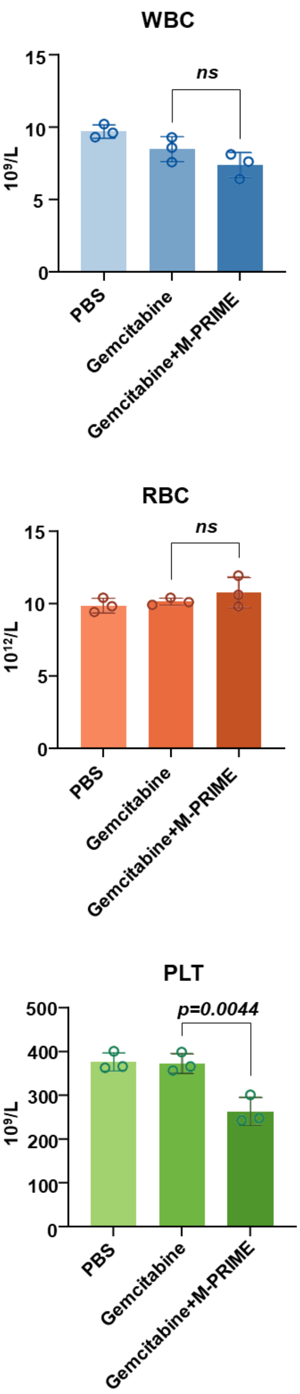


**Supplementary Fig. 26**: Exploratory haematological readouts in the orthotopic H1299-Luc lung-tumour model. Hematological parameters were measured at the experimental endpoint in mice treated with PBS, gemcitabine or gemcitabine/M-PRIME. Blue bars indicate leukocyte-related parameters, red bars erythrocyte-related parameters and green bars platelet-related parameters. Data are mean ± s.d. (n = 3 mice per group for quantitative analyses). Platelet counts were lower in the combination group than in the gemcitabine group ($p$ = 0.0044).

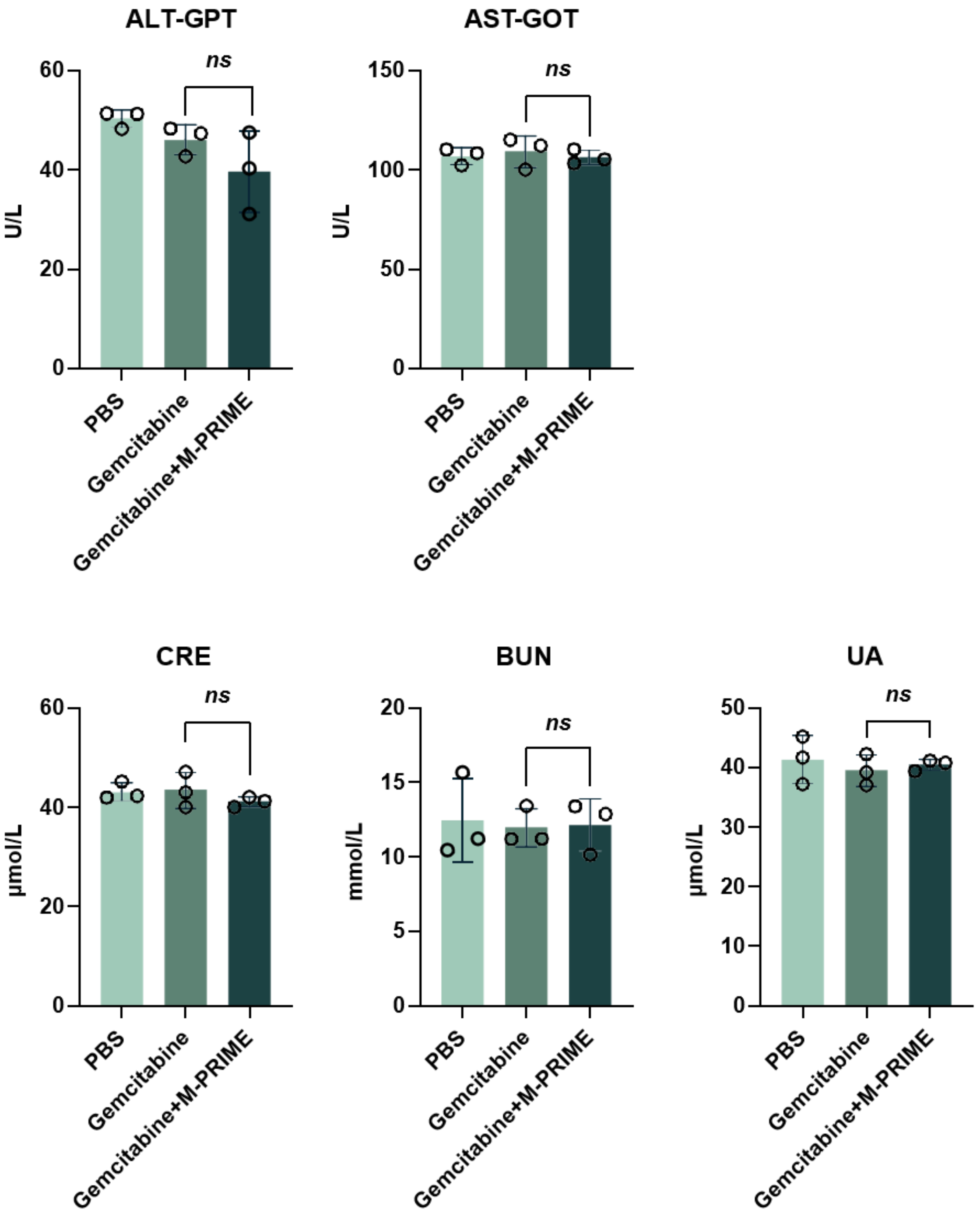


**Supplementary Fig. 27**: Serum-biochemistry safety readouts in the orthotopic H1299-Luc lung-tumour model. Serum biochemical parameters were measured at the experimental endpoint in mice treated with PBS, gemcitabine or gemcitabine/M-PRIME. The measured indicators include liver- and kidney-function-related parameters, as labelled in each panel. Data are mean ± s.d. (n = 3 mice per group for quantitative analyses).

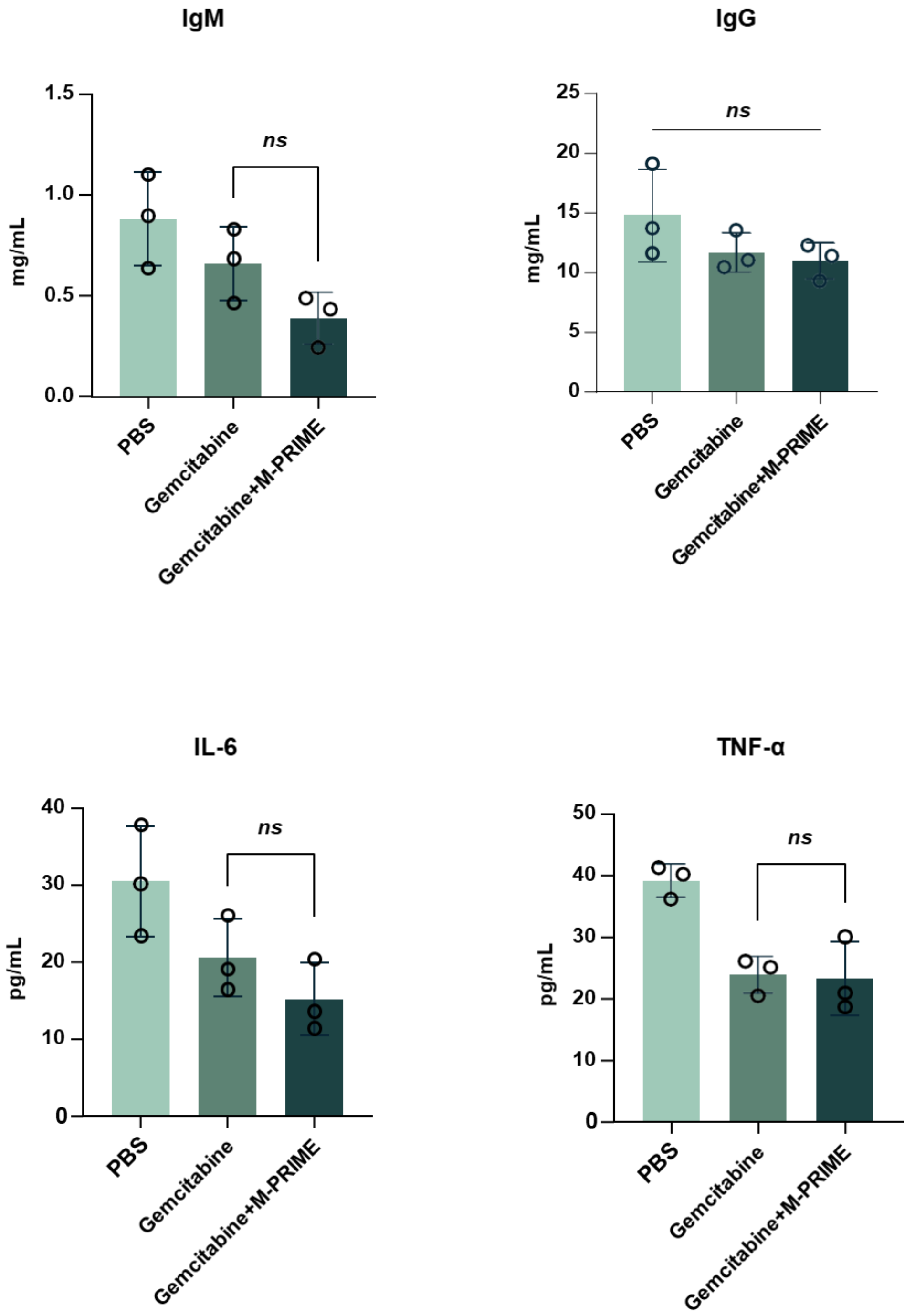


**Supplementary Fig. 28**: Plasma immune-factor readouts in the orthotopic H1299-Luc lung-tumour model. Plasma IgM, IgG, IL-6 and TNF-α were measured by ELISA at the experimental endpoint in mice treated with PBS, gemcitabine or gemcitabine/M-PRIME. Data are mean ± s.d. (n = 3 mice per group for quantitative analyses).

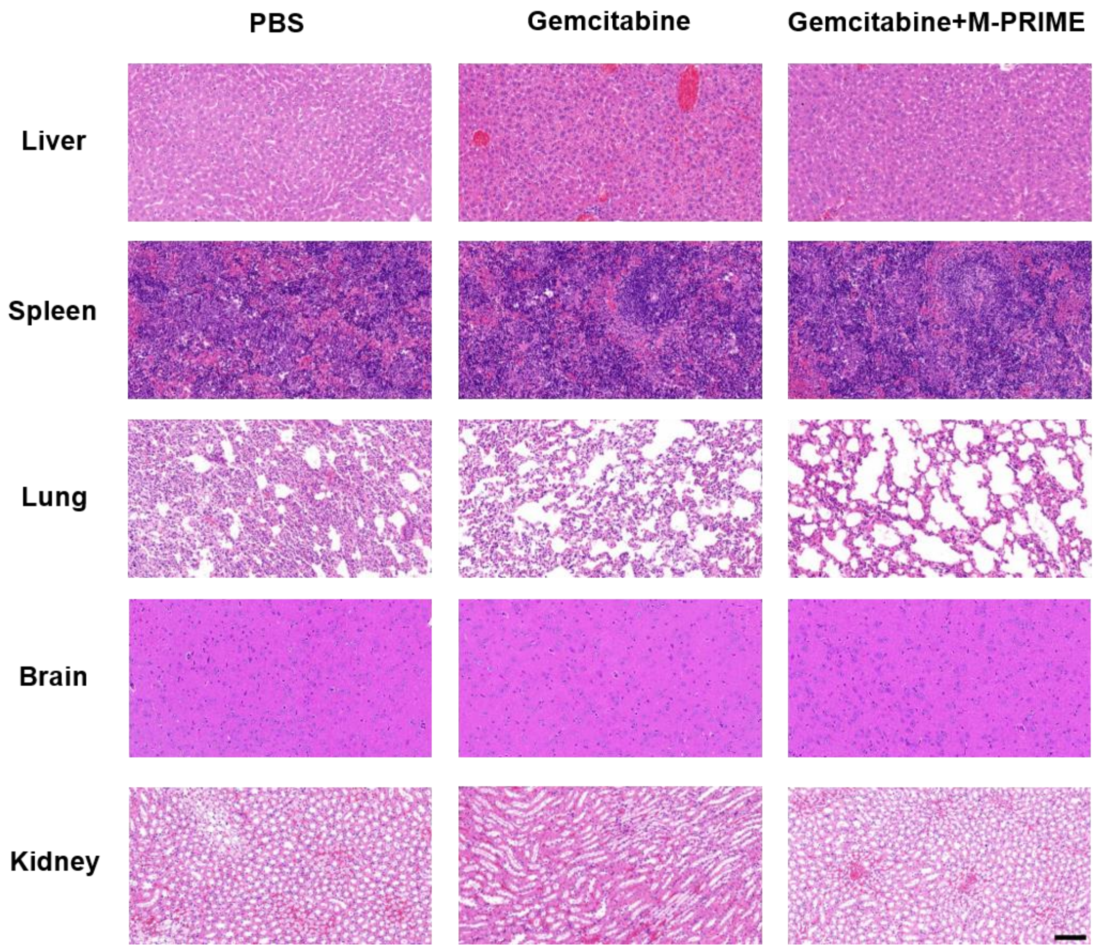


**Supplementary Fig. 29**: Major-organ histology after orthotopic lung-tumour treatment. Representative H&E-stained sections of liver, spleen, lung, brain and kidney collected from mice bearing orthotopic H1299-Luc lung tumours after treatment with PBS, gemcitabine or gemcitabine/M-PRIME. Organs were collected at the experimental endpoint, fixed, sectioned and stained with hematoxylin and eosin. Scale bar, 100 µm. Representative H&E images are shown with a scale bar of 100 µm.

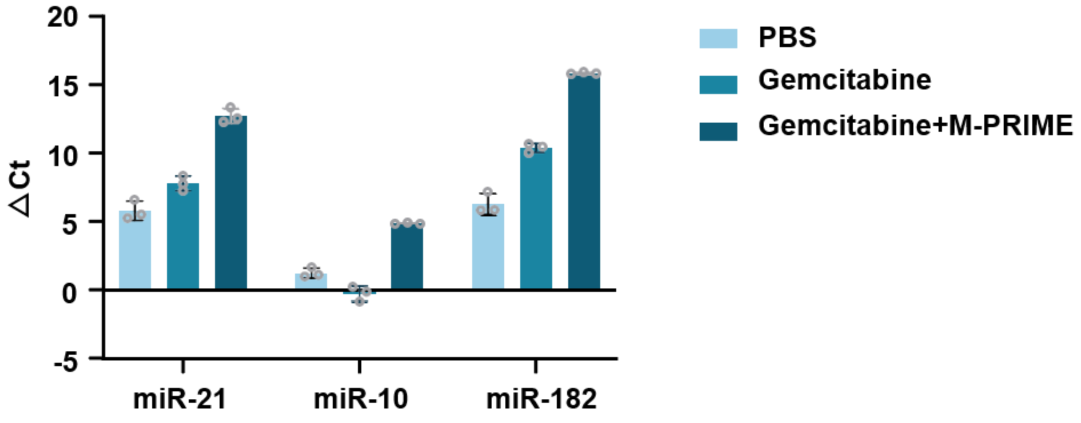

**Supplementary Fig. 30**: Raw ΔCt values for plasma miRNA RT-qPCR analysis corresponding to Fig. 7g. Plasma miR-21, miR-10 and miR-182 were measured in the MDA-MB-231-Luc hematogenous metastasis-interception model after treatment with PBS, gemcitabine or gemcitabine/M-PRIME. The y axis shows ΔCt values calculated as Ct_target miRNA - Ct_spike-in control before conversion to relative plasma miRNA abundance in Fig. 7g. Data are mean ± s.d. from plasma samples of three mice per group, each measured in technical triplicate. Each plotted point represents the mean of the three technical replicates from one mouse.

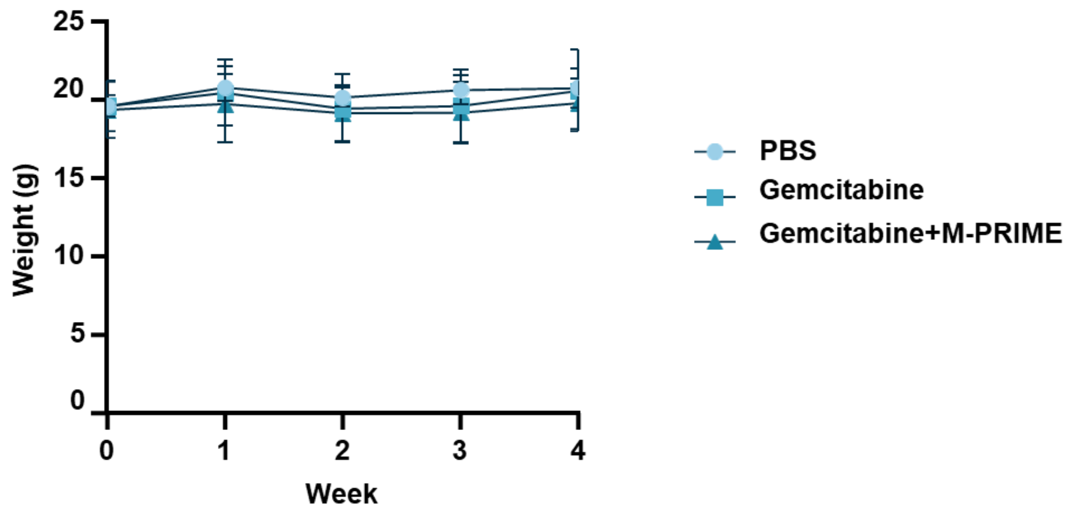


**Supplementary Fig. 31**: Body-weight monitoring in the breast-cancer hematogenous metastasis model. Weekly body-weight changes of mice treated with PBS, gemcitabine or gemcitabine/M-PRIME in the MDA-MB-231-Luc tail-vein model. Body weight was recorded once per week throughout treatment. Data are mean ± s.d. (n = 3 mice per group for quantitative analyses).

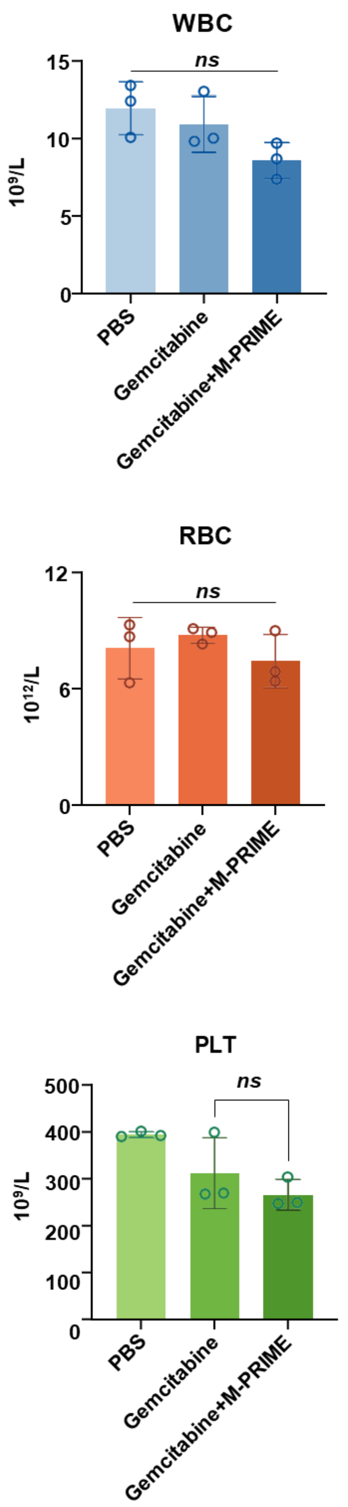


**Supplementary Fig. 32**: Hematological safety readouts in the breast-cancer hematogenous metastasis model. Hematological parameters were measured at the experimental endpoint in mice treated with PBS, gemcitabine or gemcitabine/M-PRIME. Blue bars indicate leukocyte-related parameters, red bars erythrocyte-related parameters and green bars platelet-related parameters. Data are mean ± s.d. (n = 3 mice per group for quantitative analyses).

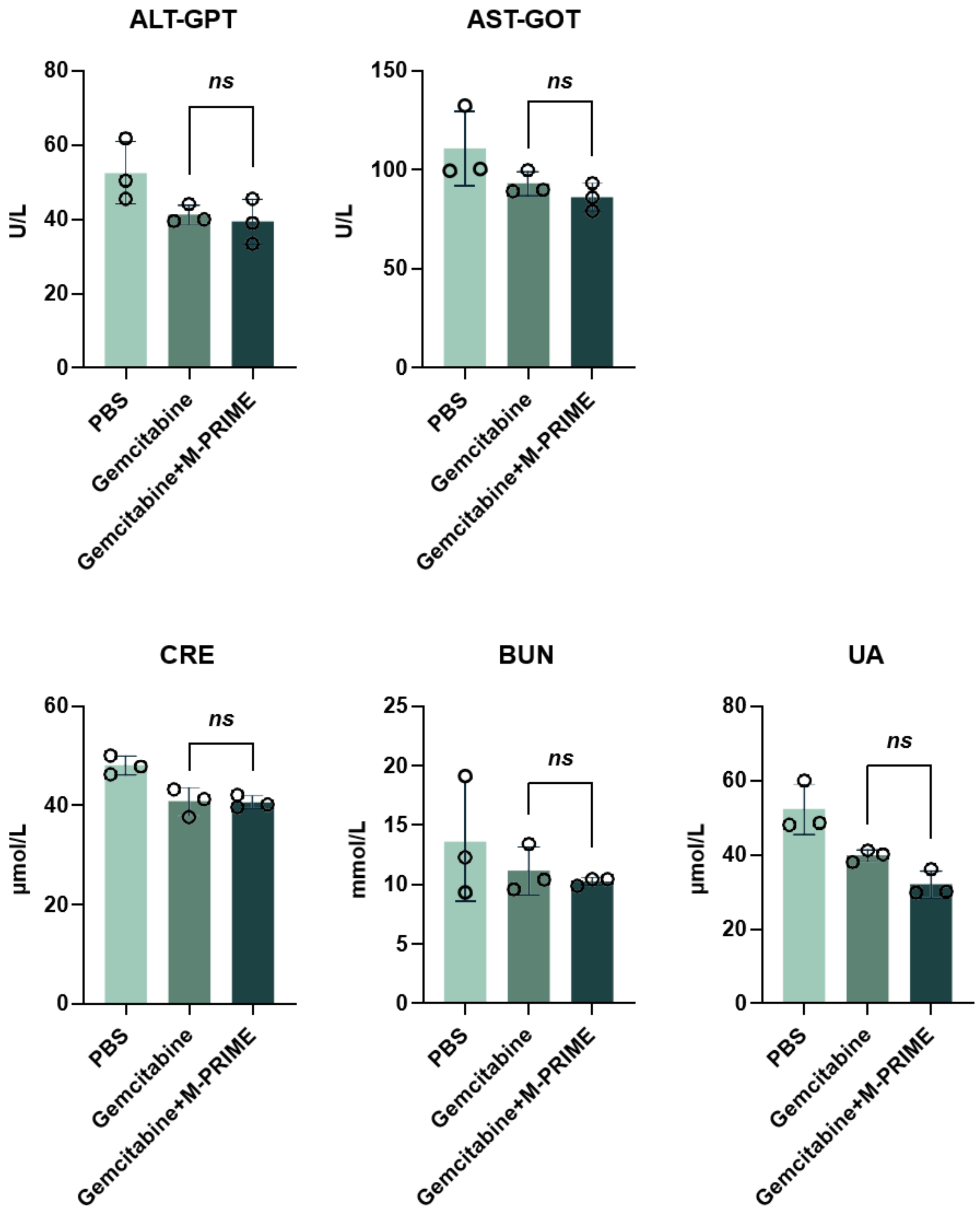


**Supplementary Fig. 33**: Serum-biochemistry safety readouts in the breast-cancer hematogenous metastasis model. Serum biochemical parameters were measured at the experimental endpoint in mice treated with PBS, gemcitabine or gemcitabine/M-PRIME. The measured indicators include liver- and kidney-function-related parameters, as labelled in each panel. Data are mean ± s.d. (n = 3 mice per group for quantitative analyses).

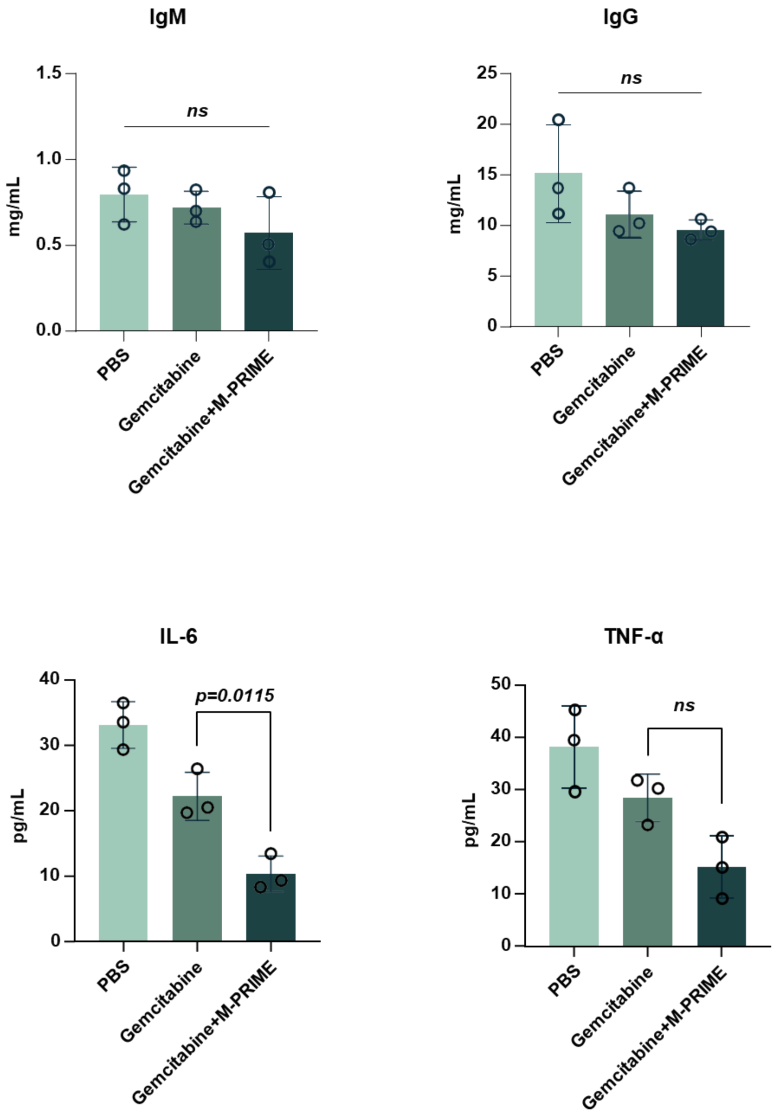


**Supplementary Fig. 34**: Plasma immune-factor readouts in the breast-cancer hematogenous metastasis model. Plasma IgM, IgG, IL-6 and TNF-α were measured by ELISA at the experimental endpoint in mice treated with PBS, gemcitabine or gemcitabine/M-PRIME. Data are mean ± s.d. (n = 3 mice per group for quantitative analyses).

**Supplementary Table 1: DNA sequences used in this study.**

| Name | Sequences (5'-3') |
|---|---|
| S1 | CACAGCGGATTCCAAACGGCACACTACGTCAGAAC AGCTTGCATCACTGGTCACCAGAGTA |
| S1-Cy5 | /Cy5/CACAGCGGATTCCAAACGGCACACTACGTCAG AACAGCTTGCATCACTGGTCACCAGAGTA |
| S2-SH | ACGAGCGAGTTGATGTGATGCAAGCTGAATGCGAG GGTCCTTTGCGAGCGCCGGCCCCCACG |
| S2-HP | ACGAGCGAGTTGATGTGATGCAAGCTGAATGCGAG GGTCCTTTCGTGGGGGCCGGCGCTCGC |
| HP-miR-21 | GCCGTTTGGAATCCGCTGTGCAGACTGATGCCCCG ACCTCAACATCAGTCGATAAGCTA |
| HP-miR-10 | GCCGTTTGGAATCCGCTGTGAGAACCGAATAACCC GACACAAATTCGGTTCTACAGGGTA |
| HP-miR-23 | GCCGTTTGGAATCCGCTGTGTGGGGATGGGCCCGC TTTAAATCCCATCCCCAGGAACCCC |
| HP-miR-182 | GCCGTTTGGAATCCGCTGTGTGGTAGAACTGGGGG GAGTGTGAGTTCTACCATTGCCAAA |
| HP-miR-92 | GCCGTTTGGAATCCGCTGTGATCGGTTGCAGAAAAT GAGCATTGCAACCGATCCCAACCT |
| HP-miR-21-BHQ3 | GCCGTTTGGAATCCGCTGTGCAGACTGATGCCCCG ACCTCAACATCAGTCT/BHQ3/GATAAGCTA |
| HP-miR-10-BHQ3 | GCCGTTTGGAATCCGCTGTGAGAACCGAATAACCC GACACAAATTCGGTTCT/BHQ3/ACAGGGTA |
| HP-miR-23-BHQ3 | GCCGTTTGGAATCCGCTGTGTGGGGATGGGCCCGC TTTAAATCCCAT/BHQ3/CCCCAGGAACCCC |
| HP-miR-182-BHQ3 | GCCGTTTGGAATCCGCTGTGTGGTAGAACTGGGGG GAGTGTGAGTTCTACCAT/BHQ3/TGCCAAA |

| Name | Sequences (5’-3’) |
| --- | --- |
| HP-miR-92-BHQ3 | GCCGTTTGGAATCCGCTGTGATCGGTTGCAGAAAATGAGCATTGCAACCGAT/BHQ3/CCCAACCT |
| S3 | TCAACTCGCTCGTAACTACACTGTGCAATACTCTGGTGACCTT |
| S4 | TCTGACGTAGTGTATGCACAGTGTAGTAAGGACCCTCGCATTT |
| S3-Apt | CTATGGGACCTCAACTCGCTCGTAACTACACTGTGCAATACTCTGGTGACCTT |
| S4-Apt | CTATGGGACCTCTGACGTAGTGTATGCACAGTGTAGTAAGGACCCTCGCATTT |
| SH-DNA | GCCGTTTGGAATCCGCTGTGAGTGGTCACCGATGAGAGAGTACTCTGGTGACCAGTGATG/SH C6/ |
| Apt-L | GGTCCCATAGGCAGTTGATCCTTTGGATACCCTGG |

Note: S1–S4 are scaffold strands used for tetrahedral assembly. S1 is the scaffold strand used to dock the programmable miRNA-recognition hairpin, whereas HP-miRNA strands contain the target-specific recognition sequences. The corresponding BHQ3-labelled HP strands are used for fluorescence-response assays. S2-SH and SH-DNA are used for construction of the RNase H–DNA catalytic module. For cholesterol-anchored PRIME, the corresponding S2, S3 and S4 strands were synthesized with a cholesterol modification at the 5′ end; their nucleotide sequences are otherwise identical to those listed in the table. S3-Apt, S4-Apt and Apt-L are used for aptamer-directed PRIME.
